# Painters in chromatin: a unified quantitative framework to systematically characterize epigenome regulation and memory

**DOI:** 10.1101/2022.03.30.486379

**Authors:** Amith Z. Abdulla, Cédric Vaillant, Daniel Jost

## Abstract

In eukaryotes, many stable and heritable phenotypes arise from the same DNA sequence, owing to epigenetic regulatory mechanisms relying on the molecular cooperativity of “reader-writer” enzymes. In this work, we focus on the fundamental, generic mechanisms behind the epigenome memory encoded by post-translational modifications of histone tails. Based on experimental knowledge, we introduce a unified modeling framework, the painter model, describing the mechanistic interplay between sequence-specific recruitment of chromatin regulators, chromatin-state-specific reader-writer processes and long-range spreading mechanisms. A systematic analysis of the model building blocks highlights the crucial impact of tridimensional chromatin organization and state-specific recruitment of enzymes on the stability of epigenomic domains and on gene expression. In particular, we show that enhanced 3D compaction of the genome and enzyme limitation facilitate the formation of ultra-stable, confined chromatin domains. The model also captures how chromatin state dynamics impact the intrinsic transcriptional properties of the region, slower kinetics leading to noisier expression. We finally apply our framework to analyze experimental data, from the propagation of *γH*2*AX* around DNA breaks in human cells to the maintenance of heterochromatin in fission yeast, illustrating how the painter model can be used to extract quantitative information on epigenomic molecular processes.

## 1 Introduction

The ability of organisms to precisely regulate gene expression is central to their development. Proper temporal and spatial expressions of genes in eukaryotes require activation of transcription during the appropriate developmental stages. In response to environmental and developmental cues, cells can adopt different gene expression patterns and differentiate into a variety of cell types. Once established, this pattern is frequently maintained over several cell divisions despite the fact that the initiating signal is no longer present or significantly weakened. This capacity of translating transient external stimuli into diverse and stable phenotypes without alteration of the genomic sequence is at the heart of the “epigenetic” regulation of gene expression [1]. Epigenetic processes are involved in the control of somatic inheritance and in the maintenance of cellular identity as well as in the transgenerational inheritance of traits by transmission via the germline [2].

A major class of mechanisms driving such epigenetic “memory” relies on chromatin-based processes regulating gene expression by the control of the local biochemical and structural properties of chromatin leading to different chromatin states more or less permissive to transcription [3]. Such control is mediated in part by biochemical modifications of the DNA or of histone tails, the so-called epigenomic marks. The mechanisms of establishment and heritability of these chromatin states, either active or repressive, are governed by similar general rules involving the combined and self-reinforcing action of specific chromatin-binding proteins that add or remove epigenomic marks [4–6].

De novo assembly first proceeds by a nucleation (“forcing”) stage via the targeting of specific enzymes at dedicated regulatory sequences by either DNA binding proteins or the RNAi-based pathways [4, 7–11]. Such nucleation elements are often composed by multiple binding sites for different DNA binding proteins that associate to their cognate sequence to stably recruit chromatin regulators. For example, in Drosophila, the Polycomb-based epigenetic repression of developmental genes relies on the targeting of the histone modifying enzymes (HMEs) PRC1 (monoubiquitination of H2AK118) and PRC2 (methylation of H3K27) complexes at specific “silencers” regions, the so-called Polycomb Response Elements, characterized by various combinations of binding sites for adaptor proteins (e.g. Gaga, Zeste, Pho) [12, 13]. Similarly, gene activation relies on the recruitment by Trithorax-group proteins (e.g. MLL2) of acetyltransferases (e.g. p300) and demethylases (e.g. UTX) at gene promoters and enhancers [14, 15].

Once initiated, the state is able to spread to the neighboring sequences and to form a stable chromatin domain [10] that can further propagate through replication and mitosis [16]. The ubiquitous ability of some chromatin regulators to be recruited by (e.g. Clr4) or to have a boosted activity in presence of (e.g. PRC2) the chromatin state they catalyze (“reading”capacity) and to spread this state to the neighbouring sequences (“writing” capacity) introduce an effective positive feedback which is believed to be a key ingredient of epigenetic maintenance [10, 16, 17].

Such “reader-writer” principle of chromatin enzymes and its impact on epigenetic memory has been already investigated using simple mathematical models of chromatin state regulation with a focus on the dynamics of histone marks mediated by HMEs [11, 18–29]. In particular, in their seminal work, Dodd *et al*. [18] suggested that the maintenance of stable, extended active or repressed chromatin domains over generations, even in absence of nucleation signals, is made possible by the reader-writer property of HMEs coupled to their capacity to spread (or write) a mark at long-range along the genome. Indeed, such interplay leads to cooperativity and allows the effective formation of a large reservoir of modified nucleosomes to serve as templates to ensure full recovery after random perturbations such as transcription-[26, 28] or replication-mediated [22, 30] histone turnover. Actually, the long-range spreading property reflects the polymeric nature of the genome that that can bring in close spatial proximity two distant loci. Many experimental and theoretical studies have indeed highlighted the correlation between spatial chromosome organisation and chromatin regulation [31–38], strengthening this hypothesis that the 3D genome impinges on chromatin states assembly and maintenance.

The reader-writer ability of HMEs coupled to long-range spreading however raises concerns about the maintenance of a stable compartmentalization of the genome [24]. Indeed, as observed in many species, the states of chromatin along the genome are linearly organised into consecutive “domains” of finite sizes, leading globally to a 1D compartmentalisation of the genome into active and inactive domains with more or less well defined inter-domain boundaries [33, 40, 41]. Because of the requirement to regulate (activation or silencing) a given part of the genome without affecting its surrounding flanks, chromatin states have thus to be specifically targeted to and stably confined inside specific genomic domains. In standard mathematical models [18, 21, 30], long-term epigenetic memory goes with great difficulties to limit the expansion of the mark, questioning mechanistically how the 1D partitioning of the genome is established and above all maintained. Several hypotheses have been proposed to address this question. Spreading may be slowed down by boundary elements [6, 19, 26, 37], usually termed “insulators”, that restrict the local spreading of a mark along the genome. These insulators are often associated with specific DNA-binding proteins (e.g. CTCF, BEAF, CP190) or actively transcribed genes found at the boundaries between antagonistic chromatin states [42–49]. Spreading may also be limited by the formation of spatial compartments [37–39, 50, 60] as genome 3D compartmentalization is strongly correlated with 1D chromatin segmentation [31, 33, 36], thus providing 3D insulation and restricting long-range effects.

Nevertheless, recent experimental studies suggest that, in vivo, the reader-writer mechanism (along with long-range spreading and insulation) might not be strong enough to self-sustain by itself an epigenetic state and that a (compartmentalized) long-term memory may still be dependent on genomic bookmarking by keeping a weak “forcing” activity at regulatory sites [10, 51–53]. Such role of nucleation signals in the maintenance of a stable chromatin state has only been partially addressed [24,27] and a general mechanistic framework of epigenomic regulation and memory integrating nucleation, reader-writer mechanisms and long-range spreading is still lacking.

In this paper, we provide such a framework and systematically investigate the formation and memory of chromatin states. In particular, our modular approach allows to carefully examine the role of sequence-specific nucleation signals, and how they interplay with reader-writer mechanisms and 3D organisation. After a deep analysis of the generic properties of each contributions, we show that such framework allows to rationalize the maintenance of confined chromatin states and to address the role of epigenome regulation in gene expression. Finally, we contextualize our quantitative approach to several concrete experimental systems, from the formation of *γ*H2AX domains around double-stranded breaks and the formation of heterochromatin domains around Transposable Elements (TE) insertion sites in mammals to the heterochromatin memory in fission yeast. The very good agreement between our simple model predictions and experimental data suggests that such multi-modal spreading model may provide a very promising framework to further investigate the link between genomic and epigenomic organisation during normal development, pathologies as well as during evolution.

## 2 MATERIALS AND METHODS

Our work consists of three main methodological parts: (i) modeling chromatin state dynamics; (ii) modeling its impact on transcription; (iii) comparing model predictions with experimental data.

### 2.1 Chromatin state dynamics

#### 2.1.1 Model

As in our previous studies [21, 22, 29], chromatin is modeled by a unidimensional array of *n* nucleosomes. As a proxy for the local chromatin state, we assume that each nucleosome can fluctuate stochastically between a finite number of epigenetic states, each state corresponding to a specific combination of histone marks. To simplify, we consider a generic two-state model between an unmodified/neutral state (U) and a modified, either active or inactive, state (M) (Fig. 1A). The switching dynamics from M to U (respectively from U to M) is controlled by the transition rate *r*_*MU*_ (resp. *r*_*UM*_).

**Figure 1.**
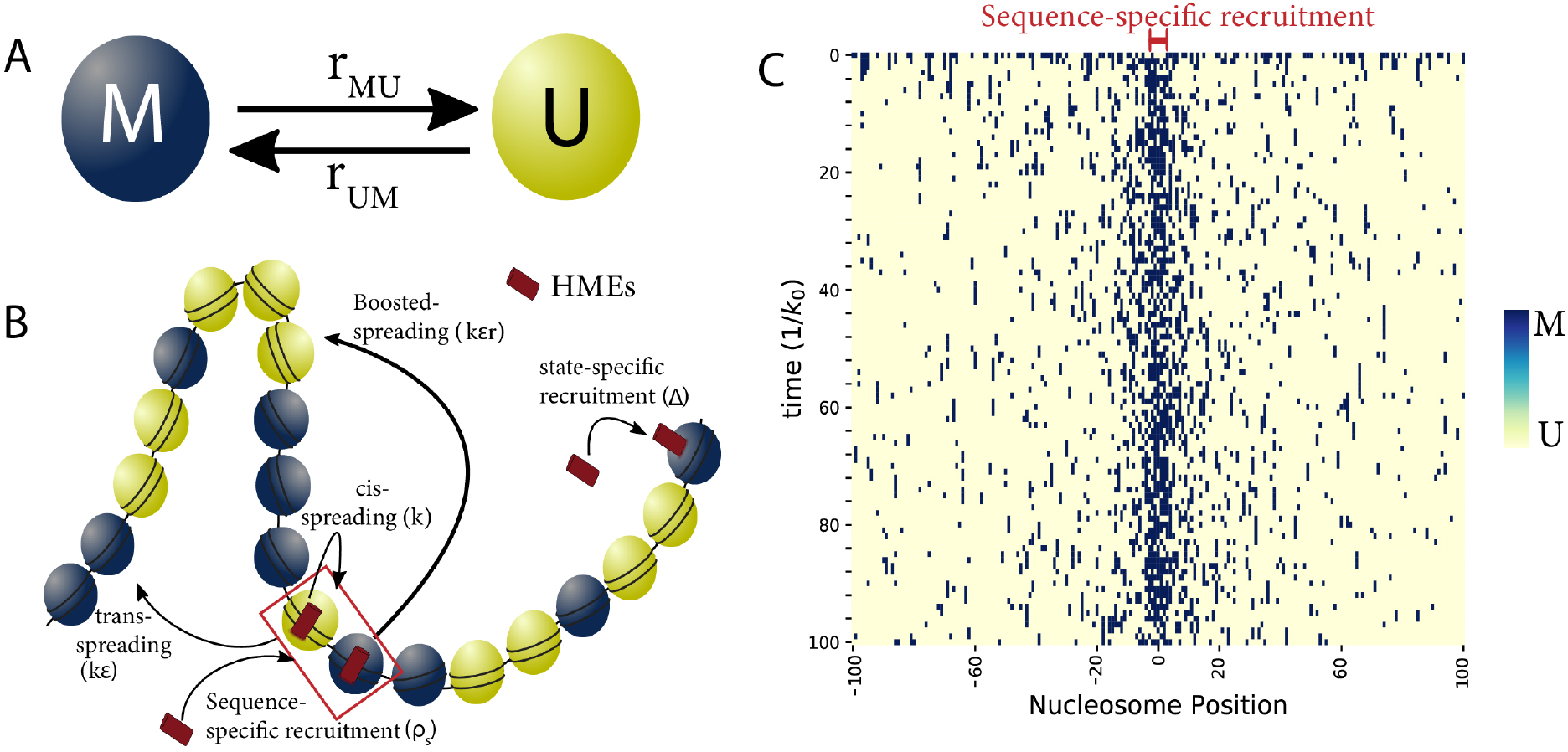
“Painter” Model: (A) Each nucleosome can be in one of the two, unmodified “U” or modified “M”, states. The switching between the two states is controlled by transition rates *r*_*UM*_, *r*_*MU*_. (B) Scheme of functionally distinct spreading mechanisms (respective terms are shown as in Eq. 1). The painter region where HMEs are recruited in a sequence-specific manner is framed in red. (C) An example of the stochastic time evolution of the epigenomic state simulated for a 201 nucleosomes-wide region in the simple painter mode.

*r*_*MU*_ integrates all the molecular processes that promote the removal of the histone modifications. This includes histone turnover [54], enzymatic removal of marks by “erasers” (such as histone deacetylases or demethylases) [55] and histone dilution between sister chromatids at replication forks [56]. In our theoretical study, for simplicity, we lump all these processes into one constant *r*_*MU*_ ≡ *k*_0_, except in Fig. 7D where we explicitly account for replication by forcing the transition from M to U with a probability 1/2 every generation time *T*_*cyc*_ = 55*/k*_0_.

*r*_*UM*_ accounts for the spreading of the state M by dedicated “writer” enzymes: HMEs are first recruited at some positions and then may “write” the M state to any neighboring U nucleosomes (Fig. 1B). We assume HMEs can be mobilized to chromatin via two independent, additive, sequence- or state-specific, pathways. The quantity of enzymes bound at position *i, ρ*_*w*_(*i*), is then given by *ρ*_*w*_(*i*) = *ρ*_*s*_(*i*) + Δ*δ*_*i,M*_ with *ρ*_*s*_(*i*) the sequence-specific contribution, *δ*_*i,M*_ = 1 if *i* is in state M, = 0 otherwise, and 0 ≤ Δ ≤ 1 is the “reader” recruitment strength accounting for the capacity of some HMEs to co-associate with the same mark they catalyze [5]. For clarity, we only consider logical distributions for *ρ*_*s*_ with *ρ*_*s*_(*i*) = 1 at given sequence-specific recruitment sites, called the “painter” regions.

Similarly, the activity *k*_*w*_(*i*) of a bound enzyme may be sequence- or state-specific: *k*_*w*_(*i*) = *k*_*s*_(*i*)(1+*rδ*_*i,M*_) with *k*_*s*_(*i*) the normal enzymatic activity and *r >* 1 a boost factor accounting for the enhancement of enzymatic activity occurring for some HMEs in the presence of the same mark they catalyze [17]. For simplicity, we assume that *k*_*s*_(*i*) ≡ *k* is homogeneous.

We consider that a bound enzyme at position *i* may spread the state M not only at nucleosome *i* (with an on-site (*in cis*) activity *k*_*w*_(*i*)) but also at longer-range (*in trans*) to any other nucleosome *j* at a rate *k*_*trans*_(*i* → *j*) = *ϵk*_*w*_(*i*)*P*_*c*_(*i, j*), proportional to the intrinsic enzyme activity *k*_*w*_(*i*), to *ϵ* (0 ≤ *ϵ* ≤ 1) a multiplicative factor accounting for a putative modulation of the writing efficiency *in trans* and *P*_*c*_(*i, j*) (≤ 1) the probability that *i* ‘communicate’ or ‘contact’ with *j* to allow spreading (see below). Altogether, the propensity for a nucleosome *i* to switch from U to M is thus given by the activities of locally bound HMEs and of HMEs bound to other nucleosomes that may interact with *i*:

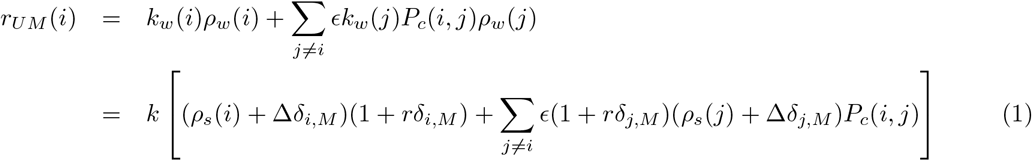

#### 2.1.2 Shape of the spreading probability

The capacity for a HME recruited at position *i* to spread a mark at long-range is encoded into *P*_*c*_(*i, j*). *P*_*c*_ may translate various physical mechanisms that allow *i* to ‘communicate’ with *j*.

A natural mechanism is to consider that *P*_*c*_ captures the frequency that *i* and *j* are in spatial proximity such that a HME bound to *i* may catalyze a reaction in *j* (Fig. 5). In this case, assuming that the 3D chromatin organization equilibrates locally faster than the epigenomic landscape, *P*_*c*_(*i, j*) can be approximated by the contact probability between *i* and *j* [60] as observed in Hi-C experiments [61]. Indeed, for short genomic distances between *i* and *j* (*<* 100kbp), theoretical and experimental studies suggest that, at this scale, 3D looping rates are ∼ sec^*−*1^ to min^*−*1^ [57, 58, 62, 63, 124] while the chromatin modification rates are often ∼ h^*−*1^ [29, 59]. In this paper, for most of our theoretical results (except in Fig. 7C,D and some cases in Fig. 5, see below), we consider a homogeneous 3D contact probability *P*_*c*_(*i, j*) = 1*/*|*i* − *j*|^*γ*^, with *γ* reflecting the average compaction level of the chromatin fiber. Experimentally, *γ* has been shown to vary in the range [0.5 − 1.5] depending on cell cycle stage, organism and/or cell fate [64–66]. In particular, we use the standard, intermediate value *γ* = 1. Note that the value of *γ* does not impact qualitatively the main conclusions of our work but may quantitatively have significant effects on the spreading process (Supplementary Fig. 1) and thus must be carefully adjusted if possible when considering specific experimental systems. In Fig. 7C,D and in Fig. 5, we also consider, for *P*_*c*_(*i, j*), contact matrices, representative of chromatin regions with (i) a central strongly-self-interacting domain of nine nucleosomes; or (2) long-range loops between the painter region (positions [−2 : 2]) and five nucleosomes at positions [48 : 52]. In both cases, *P*_*c*_(*i, j*) was computed from polymer simulations using the lattice kinetic Monte-Carlo model developed in [58] (see Supplementary Information for details). In Fig. 5, we also consider three alternative mechanisms for *P*_*c*_: (1) HMEs bound at position *i* may impact only nearest-neighbor (NN) nucleosomes, i.e. *P*_*c*_(*i, j*) = 1 if |*j* − *i*| = 1 and *P*_*c*_(*i, j*) = 0 otherwise; (ii) HMEs recruited at *i*, while unbinding and diffusing, are more likely to rebind in the 3D vicinity, leading to a concentration gradient around *i* and to an effective *P*_*c*_(*i, j*) = 1*/*|*i* − *j*|^0.5^ (see Supplementary Information); (3) HMEs bound at *i* may spread a mark to a distal nucleosome *j* only if they are placed in very closed proximity by a loop extruding factor (like cohesin or condensin) [67–70] translocating along the chromatin, leading to *P*_*c*_(*i, j*) = exp(−|*j* − *i*|*/s*_0_) (see Supplementary information and Supplementary Fig. 2).

#### 2.1.3 Stochastic simulations

For a given set of parameters, the stochastic dynamics of the system is simulated using the standard Gillespie algorithm [71] implemented on Python (can be downloaded at https://github.com/physical-biology-of-chromatin/Painter-Model). In all our work, we simulated the dynamics of *n* = 201 nucleosomes corresponding to ∼ 40kbp-long genomic region. Starting from a random initial macro-state (i.e random choice between U or M states for each nucleosome), the system relaxes to a steady state. Each simulation corresponds to a “single cell” trajectory of the local epigenetic state (Fig. 1C). Unless specified (Fig. 6, 7), characterization of the system was done at steady-state. In each condition, 500 different trajectories were simulated. Times are given in 1*/k*_0_ unit that characterizes the typical turnover time of histone marks and is ∼ *h* [29, 54].

#### 2.1.4 Analytical solutions

*In the case where state-specific terms are negligible (r* = Δ = 0, painter mode), the steady-state probability *P* (*M*_*i*_) of being in modified state M at any position *i* is simply given by :

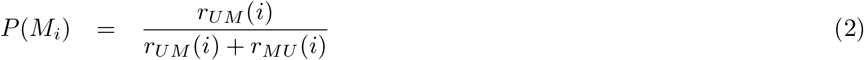

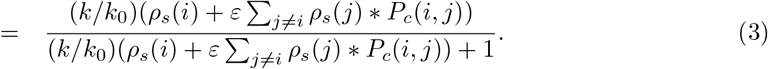

In the case where only state-specific recruitment is negligible (Δ = 0) and where the sequence-specific recruitment is localized in a finite painter region, by doing a mean-field approximation, the average probability of the M-state inside the painter region 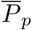 is given by analytically solving (see Supplementary Information and Supplementary Figure 3):

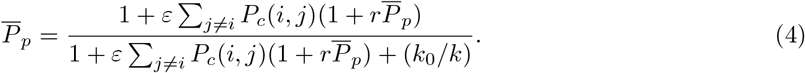

Outside the painter region, *P* (*M*_*i*_) is then given by replacing *ε* by 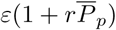 in Eq. 3.

#### 2.1.5 Enzyme limitation

In the model described above, we assumed that the concentration of HME is large enough to not have to consider the depletion of the pool of freely-diffusing HMEs (that impacts the sequence- and state-specific recruitment strength) by bound HMEs. In Fig. 7, we consider a scenario where the number of enzymes is actually limited. Assuming fast binding-unbinding kinetics for the enzymes, we can show (see Supplementary Information and Supplementary Fig. 4) that such system is equivalent to our simple model (Eq. 1) but by replacing the state-dependent parameter Δ by *P* (*E*) the probability that a HME is bound to a modified nucleosome. *P* (*E*) explicitly depends on 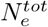 the total number of HMEs, *k*_*b*_ and *k*_*u*_ the binding and unbinding rates (see Eq. 7 in Supplementary Information). In the limit of a non-limiting number of enzymes 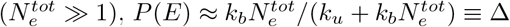. In Fig. 7, we fix *k*_*b*_*/k*_*u*_ = 1.

#### 2.1.6 Contact domains

The spreading probability *P*_*c*_(*i, j*) represents how frequent pairs of nucleosomes are ‘in (effective) contact’. Therefore, the genomic region is not only a unidimensional string of beads with only NN links but a network of interactions. In order to capture at the single-cell level, the propensity of nucleosome in the M-state to be ‘connected’, we randomly generated for every configuration an undirected graph (*M*_*net*_) for the corresponding set of M-states (Supplementary Fig. 5). More precisely, for all pairs (*i, j*) of nucleosomes in this set, an edge between them is inserted in *M*_*net*_ with a probability *P*_*c*_(*i, j*). Then, we infer the disconnected subgraphs of *M*_*net*_ that would represent the different clusters of connected M-state nucleosomes. Finally, for each node/nucleosome *i*, we compute the size of the contact domain it belongs to.

In the reader-writer mode, a M-state nucleosome may influence a U-state nucleosome via *P*_*c*_. Thus, to behave coherently, a genomic region may not necessarily need to have all nucleosomes in the M-state, but rather that all M-states belong to the same graph and that all U-states can be reached via *P*_*c*_(*i, j*) and are connected to such network. To quantify this, for each configuration, we consider the largest M-state subgraph *M*_*net*_∗ in *M*_*net*_. For each U-state nucleosome *i*, random links with any M-state nucleosome *j* of *M*_*net*_∗ are generated with probability *P*_*c*_(*i, j*). *i* is added to *M*_*net*_∗ if it has been linked to at least two M-state nodes of *M*_*net*_∗ (see Supplementary Fig. 5, Fig. 5C). The size of such extended contact domain can thus be considered analogous to a percolation parameter [72], indicative of the extent of M-state spreading in the region.

#### 2.1.7 Nucleosome state correlations

To compute the spatial correlations between nucleosome states, we divide the genomic region into 40 bins of 5 nucleosomes. In one configuration, these bins are thus characterized by a M-state ranging from 0 to 5. Nucleosome state correlation matrix shown in Fig. 2, 3, 4C represents the Pearson correlation between the M-states computed for each pair of nucleosomes over all the configurations extracted from the simulations.

**Figure 2.**
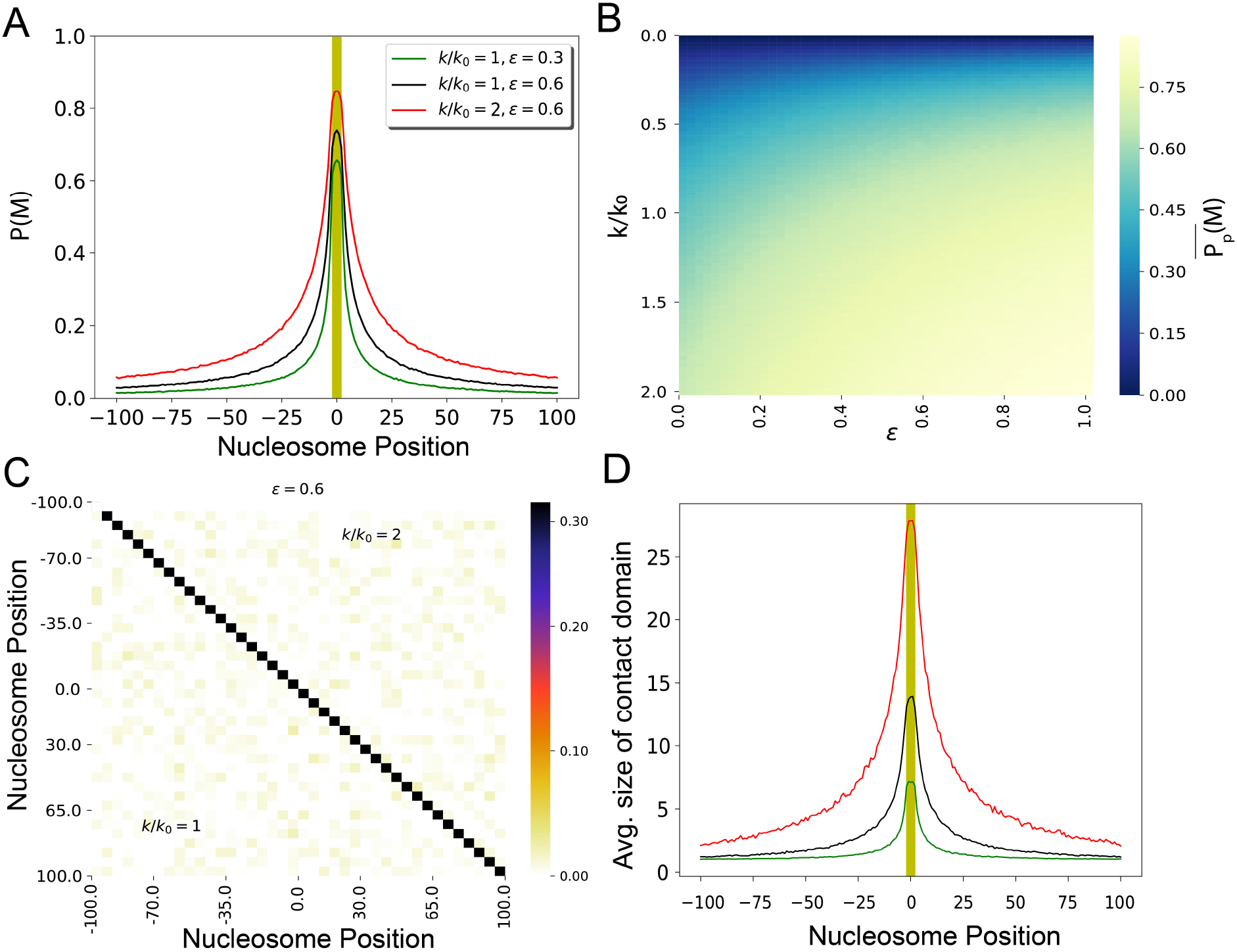
Sequence-specific recruitment - Painter mode : (A) Virtual Chip-seq profiles of the M-state. Probability *P* (*M*) to be modified as a function of the position around the 5 nucleosomes-long region where HMEs are recruited (yellow region) for different spreading efficiency *ε* and painter activity *k/k*_0_. (B) Average probability 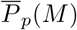 of the M-state within the recruitment region as a function of *ε* and *k/k*_0_. (C) Spatial correlation matrix between the nucleosome state at two positions (see Materials and Methods) for *ε* = 0.6 and *k/k*_0_ = 1 (lower part) or = 2 (upper part). (D) Average M-state contact domain size as a function of the nucleosome position for different *ε* and *k/k*_0_ values.

**Figure 3.**
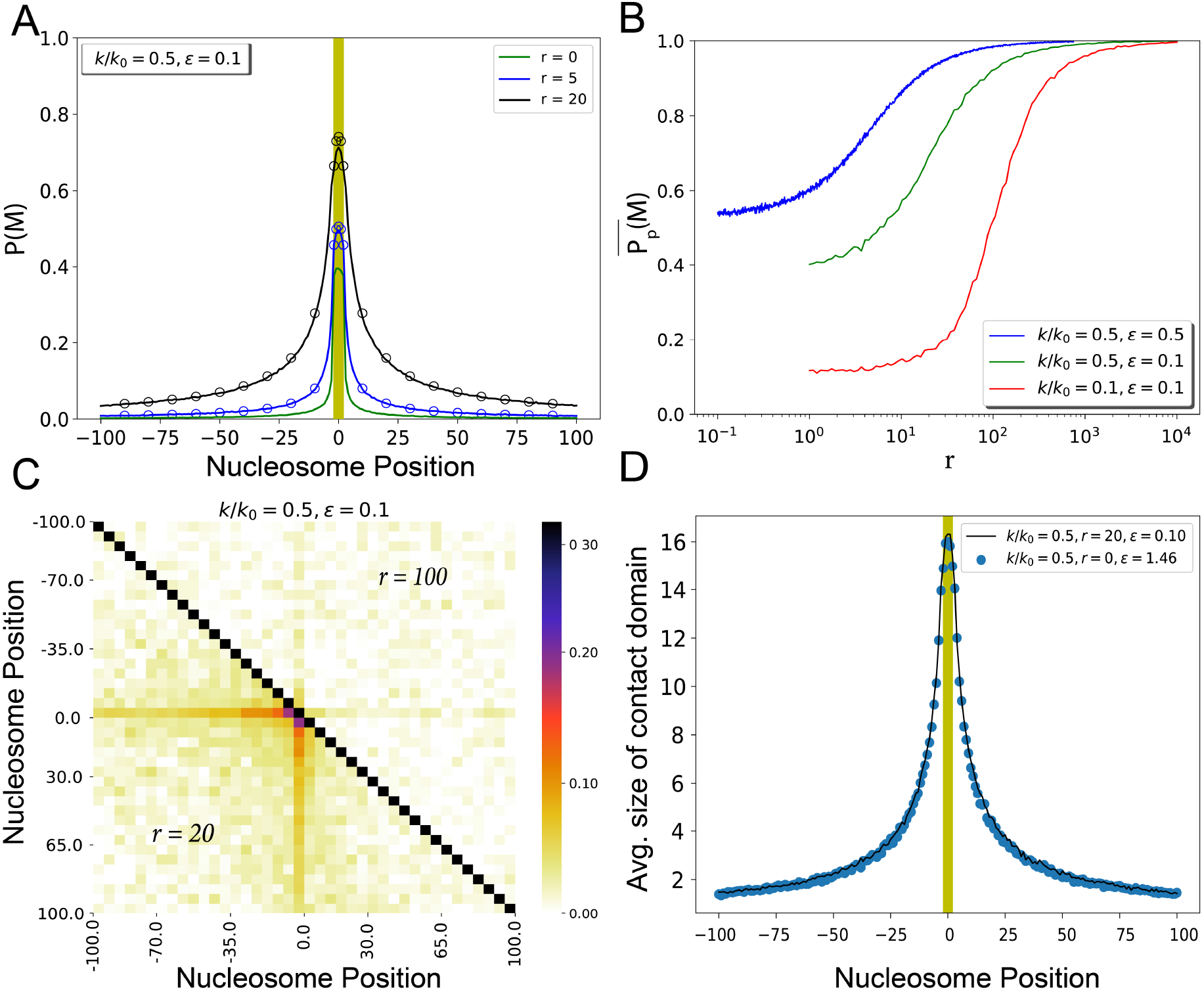
State-specific enzyme activity - boosted painter mode : (A) Virtual Chip-seq profiles of the M-state. *P* (*M*) as a function of the position around the 5 nucleosomes-long region where HMEs are recruited (yellow region) for different values of the boost factor *r*. The scatter points are the analytical solution of the model. (B) 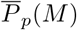 within the recruitment region as a function of *r*. (C) Spatial correlation matrix between the nucleosome state at two positions for *ε* = 0.1, *k/k*_0_ = 0.5 and *r* = 20 (lower part) or = 100 (upper part). (D) Average M-state contact domain size as a function of the nucleosome position with boost *r* = 20 (full line). Dots correspond to a simple painter mode with an adjusted *ε* value to reproduce the *P* (*M*) profile.

**Figure 4.**
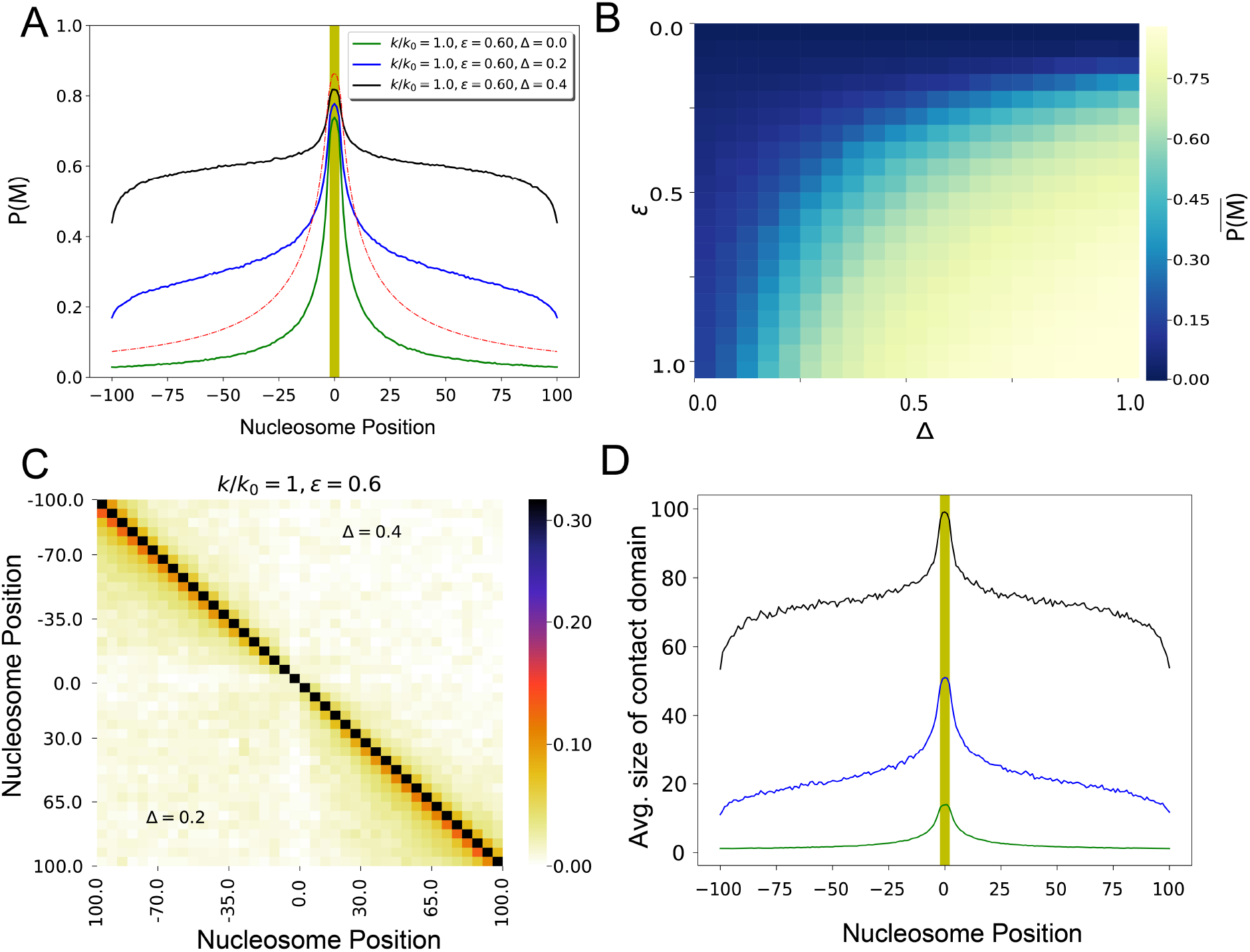
State-specific recruitment of enzyme - reader-writer mode: (A) Virtual Chip-seq profiles of the M-state. *P* (*M*) as a function of the position around the 5 nucleosomes-long region where HMEs are recruited (sequence specific) for different values of reader recruitment strength Δ (at the critical point Δ = 0.2 and above = 0.4). (B) Average value 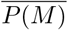 of *P* (*M*) inside the whole genomic region as a function of *ε* and Δ (*k/k*_0_ = 1). (C) Spatial correlation matrix between the nucleosome state at two positions for *ε* = 0.6, *k/k*_0_ = 1 and Δ = 0.2 (lower part) or = 0.4 (upper part). (D) Average M-state contact domain size as a function of the nucleosome position for different Δ values. Colors as in (A).

### 2.2 Transcription dynamics

Assimilating the modified state M to a silencing state, we assume that the instantaneous transcription rate *α* of a gene localized inside the region of interest is negatively impacted by the current proportion of M-state nucleosomes inside the promoter region. More precisely, we use the cooperative switch model of transcription (Fig. 8A), proposed by Zerihun *et al*. [22].

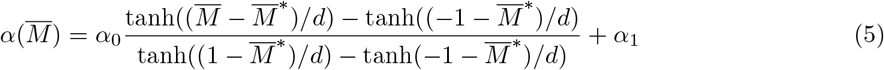

where *α*_0_ + *α*_1_ is the maximal transcription rate, *α*_1_ is the minimal, leaky, rate, 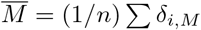 the current proportion of modified state in a *n* nucleosome-wide promoter region around, 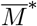 the critical value of modification above which the gene is repressed and *d* defines the sharpness of the transition between activation and repression.

Gene expression dynamics is simulated along with the chromatin state dynamics within the same Gillespie simulations by considering, in addition to {*r*_*UM*_ (*i*), *r*_*MU*_ (*i*)}, the transcription of mRNA molecules at rate *α* (Eq. 5), their degradation at rate *β*, their translation into proteins at rate Γ, the degradation of proteins at rate Ω and the maturation of proteins at rate *μ* [73]. In Fig. 8, we fix 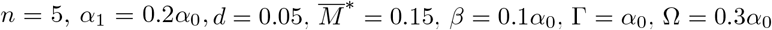 and *μ* = 0.1*α*_0_.

### 2.3 Analysis of experimental data

In Fig. 9A, the average experimental profile *S*(*i*) of H2A.X phosphorylation in human cells around double strand breaks is extracted from Sup. Fig. 1k (right) of Arnould *et al*. [70]. We fit the experimental data by the chromatin state model in the painter mode (*r* = Δ = 0) with one painter (ATM kinase) at position 0 (see Suppl. Fig. 1k, left of [70]) for a ±1Mbp-region around the break by using Eq. 3 and assuming that *S*(*i*) = *AP* (*M*_*i*_) + *B* with *A* and *B* two constants. Parameter inference is done by minimizing the L2-distance between predictions and experiments.

Similarly, in Fig. 9B, the average density of H3K9me3 modifications flanking a transposable element in mESC cells is obtained from Robollo *et al*. [74]. Fit to experimental data using Eq. 3 is performed for a 10kbp-region with a centered painter region of size 7 nucleosomes corresponding to 1kbp transposable element. Parameter inference is done by minimizing the L2-distance between predictions and experiments.

In Fig. 9C the distribution of fluorescence of the reporter gene as a function of time and the corresponding fractions of “OFF” cells are extracted from Ragunathan *et al*. [10] for different strains. Modeling of the system is performed using the chromatin state+transcription model (see above) with 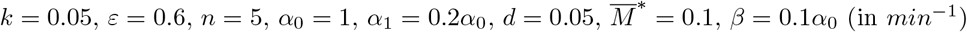 and the same parameters for protein dynamics as above. A scaling is applied to convert the number of proteins *p* into a fluorescence level *F* (*F* = *p* + *C* with *C* = 380). The fraction of OFF cells is defined by the fraction of cells whose fluorescence is below 530 as defined experimentally [10]. The extracted experimental fluorescence at 6 hours is not aligned with the fluorescence peak observed at other time points. Fit to experimental data is performed by minimizing 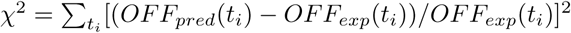, where *OFF*_*pred*_(*t*_*i*_) is the predicted fraction of OFF cells at time *t*_*i*_ and *OFF*_*exp*_(*t*_*i*_) the corresponding experimental observation, using a simple grid-search algorithm to scan parameters (see Supplementary Fig. 6).

## 3 RESULTS

### 3.1 A generic model of epigenomic regulation

To investigate in detail the main mechanisms driving epigenomic regulation, we develop a simple generic two-state model (Fig. 1A) where the local chromatin state can fluctuate between an unmodified, neutral state U and a modified state M that may correspond to an active (e.g. H3K4me3 or H3K27ac) or repressive (e.g. H3K27me3 or H3K9me2/3) state. We model chromatin as a unidimensional string of nucleosomes, each nucleosome being in state U or M. We consider a ∼40-kbp-long genomic domain (201 nucleosomes), a size that typically encompasses epigenetically-regulated regions like transposable elements [75], gene promoters and enhancers, the Mating type locus or subtelomeres in yeast [76].

The stochastic dynamics of the state is driven by histone-modifying enzymes (HMEs) that deposit (“writers”, transition from U to M) or remove (“erasers”, from M to U) specific histone modifications [5,10,16]. For example, PRC2 via its methyltransferase subunits EZH1 or EZH2 catalyzes the methylation of H3K27 [2, 77]. Our model integrates the key mechanisms acting on HME recruitment or activity, focusing on “writer” enzymes and lumping all the processes participating in histone mark removal into one effective turnover rate *k*_0_ (see Materials and Methods for details). Briefly (Fig. 1B), we have delineated the chromatin association of HMEs into sequence- and state-specific contributions: (a) writers, in association with DNA-binding proteins, may localize around specific genomic location with probability *ρ*_*s*_, or (b) HME may be recruited to M-state nucleosomes with efficiency Δ. Recruited HMEs at position *i* can then write the epigenetic mark on-site with a rate *k* (action *in cis*) but may also spread it to a distal nucleosome *j* with a rate (*εk*)*P*_*c*_(*i, j*) (action *in trans*) [15], where *P*_*c*_(*i, j*) accounts for the capacity of two nucleosomes to interact and may depend on the exact spreading mechanism (see Materials and Methods). In the following, we consider that bound HMEs may catalyze reaction in their 3D neighborhood, as evidenced experimentally for several epigenetic systems including H3K27 methylation [15, 83], H3K9 methylation [78], *γ*H2AX phosphorylation around DNA double strand breaks [79] and *P*_*c*_(*i, j*) is thus defined as the average 3D contact frequency between two genomic regions. HME enzymatic activity may be also boosted by allostery by a factor *r* if bound to a M-state nucleosome [17, 77, 80]. State-specific effects (recruitment and allosteric boost) are the so-called reader-writer mechanisms that are thought to be crucial for the establishment and maintenance of many epigenomic states [16].

In the following sections, we will systematically dissect the role of sequence-specific recruitment and of these reader-writer processes in epigenetic regulation using extensive simulations of our generic two-state model.

### 3.2 Spreading by sequence-dependent recruitment of enzymes: the “painter” mode

We first investigate the contribution of a simple mode of spreading where enzymes bind only to specific genomic locations and then may spread epigenetic marks in the 3D vicinity, in absence of reader-writer mechanisms. In this case, HMEs can be considered as “painters” that sit at specific places along the genome and “paint” the surrounding chromatin with a given modification. As sequence-dependent recruitment of enzymes is essential for *de novo* establishment of epigenetic domains [13, 81] and as many writers do not carry a “reader” subunit (that may drive state-dependent effects), such mode of regulation, hereafter called the “painter” mode, is likely to be a major way of epigenetic spreading.

To illustrate this model, we consider a simple case where painters are only recruited to a single 1kbp-long locus located in the middle of the 40kbp-long region (yellow area in Fig. 2A). In this situation, the probability *P* (*M*) to have a modified nucleosome at a given position can be derived analytically (see Eq. 3 in Materials and Methods) and only depends on the ratio (*k/k*_0_) between the cis-spreading rate *k* and the turnover rate *k*_0_ and on the ratio *ε* between the trans- and cis-activities (Fig. 2A,B). For all positions, we observe that *P* (*M*) is a gradually increasing function of both parameters (Fig. 2B, Supplementary Fig. 7). In the limit of very low trans-efficiency (*ε* ≈ 0), modified states are essentially confined within the recruitment region due to the remaining on-site writing activity. For larger *ε* values, *P* (*M*) is peaked at the painter region and decays at large genomic distances. The stronger *ε* and *k/k*_0_ the wider the peak in *P* (*M*) profiles and thus longer the range of spreading of the M-state (Fig. 2A).

Actually, the observed decay of *P* (*M*) outside the recruitment zone is translating the decay of contact probability *P*_*c*_ with the painters bound to recruitment zone. More generally, in the limit of low contact probability or low spreading efficiency ((*k/k*_0_)*P*_*c*_*ε* ≪ 1), *P* (*M*) is directly proportional to the spreading probability *P*_*c*_ (see Eq. 3) which, in our case, is taken to be ∝ 1*/s*^*γ*^ with *s* the genomic distance to the painter region and *γ* = 1 (see also Supplementary Fig. 1). This illustrates the direct relationship between the local 3D organization and the profiles of epigenomic marks around recruitment sites [82].

In the painter mode, the local chromatin state only depends on the position of the bound painters and does not feedback on the recruitment of HMEs or on their activity. This lack of cooperativity leads to the absence of correlation between the nucleosome states at two different positions along the region (Fig. 2C). To quantify the efficiency of the spreading mechanism and its capacity, at the single-cell level, to form more or less expanded, coherent M-state domain, we estimate how nucleosomes in the M-state are effectively ‘colocalized’ in space. For each configuration and each M-state nucleosome, we compute the number of other M-state nucleosomes present in the same 3D contact domain (see Materials and Methods). Fig. 2D shows that, in average, except around the painter region, M-state nucleosomes are isolated from each other and only part of small domains.

### 3.3 State-dependent enzymatic activity: the “boosted-painter” mode

In some systems such as Polycomb regulation (associated with the H3K27me3 modified state), it has been observed that the activity of the writers can be “boosted” by the presence of pre-existing modifications [17, 77, 80, 83, 84]. For example, the binding of the Polycomb writer PRC2 to H3K27me3 marked nucleosomes, triggers a boost in methyltransferase activity of its subunit EZH2 by allostery, via its other subunit EED. In our theoretical framework, this effect can be formalized by an increase in the writing rate of HMEs bound to a modified nucleosome via a multiplicative factor *r >* 1 (see Materials and Methods). For example, for mammalian PRC2, in vitro experiments suggested an allosteric boost of *r* ≲ 10 fold in presence of H3K27me3 peptides [17, 84].

In Fig. 3, we characterize the impact of this boost on the simple painter mode described above (stably bound writers at a specific 1kbp-long region), still neglecting state-specific recruitment of HMEs (Δ = 0). As shown in Fig. 3A, a weak *ε* value that essentially confines the M-state to the painter region in the simple painter mode (*r* = 0, green line), can be compensated by a strong boost term (*r* = 20, black line) with a significant overall enhancement of *P* (*M*): the presence of modified nucleosomes in the painter region due to the “on site”, cis-activity of painters (controlled by *k/k*_0_) boosts globally the cis- and trans-spreading capacity of bound writers (via *r*) and favors further spreading inside and outside the painter region.

To get a better understanding of this “boosted-painter” mode, we first focus on the painter region by computing 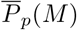, the average value of *P* (*M*) inside this region (Fig. 3B). As expected, 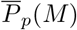, is an increasing function of *k/k*_0_ and *r*. However, contrary to the simple painter mode (Supplementary Fig. 7B), the transition from low to high M-state is a much sharper sigmoid function. Such switch-like behavior suggests a phase transition [27] inside the painter region that reflects the cooperative dynamics between writer-bound nucleosomes due to the state-dependent boost. The sharpness and position of the transition depends on *k/k*_0_ and *ε*. High values lead (i) to smoother transition as the simple painter mode is significant enough to buffer the boost effect and (ii) to lower critical *r*-values as less boost is required to get high enzymatic activity.

In this spreading mode, the chromatin states of nucleosomes localized outside the painter region are now dynamically coupled to the ones inside, leading to positive spatial correlations that are maximal for *r*-values around the critical boost (Fig. 3C). Such coupling makes an exact analytical treatment of *P* (*M*) intractable. However, using a mean-field approximation inside the painter region, we can derive an expression for 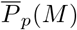 (see Materials and Methods and Supplementary Information). Outside the painter region, *P* (*M*) is thus just as in the simple painter mode but for a boosted trans-activity 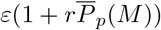 (blue and black circles in Fig. 3A). Hence, regarding the profile of epigenomic mark and of coherent 3D contact domain size (Fig. 3D), the boosted painter mode is indistinguishable from a simple painter mode with a greater “on site” and “off site” spreading rates (Blue dots, Fig. 3D). Only the presence of spatial correlations of nucleosome states with the recruitment region (Fig. 3C) might be a clear signature of this mode of propagation.

### 3.4 State-specific recruitment of enzymes: the “reader-writer” mode

Some writer enzymes have been shown to be recruited in a state-dependent manner to chromatin, having the ability to “read” (i.e. to be recruited at) a particular histone tail modification and “write” the same on another nucleosome in the 3D vicinity [2, 3, 5, 10]. For example, the methyltransferase Clr4 (associated with heterochromatin formation and H3K9me2/3 modifications in fission yeast) contains a chromodomain that may trigger its recruitment by H3K9me2/3 [10]. In our framework, we introduce this “reader-writer” mode by assuming that the probability of finding a HME bound at a M-state nucleosome is enhanced by a factor Δ (see Materials and Methods). In the following part of this section, we focus on this effect coupled to the simple painter mode neglecting possible state-specific boost (*r* = 0).

In that case, thanks to the long-range action of M-state bound writers, the state dynamics of every nucleosome is coupled to the states of all the other nucleosomes which is well illustrated in Fig. 4C by the global increase in spatial correlations. Such reader-writer mode introduces a positive feedback in the global M-state dynamics which has been shown to promote the formation and inheritance of extended, stable M state domains [18, 32]. As shown in Fig. 4A, when combined with the simple painter mode, state-specific recruitment strongly modifies the spreading pattern by notably increasing M-state occurrence away from painter region. In particular, we observe heavy tails for *P* (*M*) (black and blue lines in Fig. 4A) that qualitatively differs from the typical 1*/s*^*γ*^ contact decay observed for the sole painter mode (red dotted curve in Fig. 4A). Such signature (heavy tails, deviation from simple painter model) in experimental profiles of epigenomic marks may thus be suggestive of a dominant reader-writer mechanism. In this regime, we also observe that the reader-writer process facilitates the formation of expanded 3D M-state contact domains away from the sequence-dependent recruitment region (fig. 4D) unlike the simple and boosted painter modes.

To quantify the resulting global increase in *P* (*M*), we systematically compute 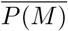, the average value of *P* (*M*) inside the whole genomic region (Fig. 4B). For a given trans-spreading activity (*ε*), we observe a sharp transition when state-specific recruitment efficiency (Δ) augments: from a pure simple painter profile localized around the painter region 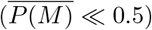 to a globally modified state 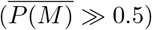. As in the boosted-painter mode, this also suggests a phase transition but here that reflects the cooperative dynamics between *all* nucleosomes of the region (see also next section). Around the critical Δ-value (e.g. Δ_*c*_ ≈ 0.2 for *k/k*_0_ = 1 and *ε* = 0.6), fluctuations in *P* (*M*) are maximal and lead to high inter-nucleosome correlations (Fig. 4C and Supplementary Fig.8).

### 3.5 The spreading probability drives the percolation of the chromatin landscape

A key element of the different writing modes investigated above is the capacity of recruited HMEs to spread an epigenetic signal [2, 3, 5, 10]. This property depends on the spreading probability *P*_*c*_(*i, j*) that captures the ability of two nucleosomes to interact. In the previous sections, we assumed that *P*_*c*_(*i, j*) = 1*/*|*j* − *i*|^*γ*^ with *γ* = 1, a generic scaling law accounting for a mechanism of spreading via 3D contacts as already observed for some HMEs like Polycomb group proteins (H3K27 methylation) [15], Clr4 methyltransferase (H3K9) [10], ATM kinase (*γ*H2AX) [70] and characteristic of the polymeric nature of chromatin [92]. In this section, we explore the role of the shape of *P*_*c*_(*i, j*) in epigenomic regulation.

This shape may depend on the specific spreading mechanism and on the experimental system or genomic region under study. Here, we consider five alternative forms for *P*_*c*_(*i, j*) (see Materials and Methods). (1) Two that still correspond to a 3D contact spreading process but with contextualized, heterogeneous *P*_*c*_(*i, j*) (Supplementary Fig. 9A): one for a small 3D compact domain localized around the recruitment zone and another for a region with a strong loop between the painter area and a locus 10kbp away, mimicking, for example, a repressed MAT locus in yeasts and a promoter-enhancer loop in mammals, respectively. (2) A shape accounting for a spreading to only nearest-neighbor (NN) nucleosomes that might be relevant in scenarios where an epigenomic state propagates unidimensionally along the chromatin [24, 91, 115]. (3) A form compatible with an effective contact spreading mechanism where HMEs recruited at a given position may diffuse in 3D to nearby positions and thus impact the state of distal nucleosomes (*P*_*c*_(*i, j*) = 1*/*|*j* − *i*|^0.5^). (4) A scenario where two nucleosomes may influence each other only if they have been placed in very close proximity by loop extruding factors (eg, cohesin or condensin) that are molecular motors translocating along the chromatin and implicated in TAD formation in mammals [67–70] 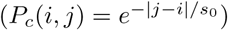.

In the simple painter mode (Fig. 5A, Δ = 0), spreading from the central recruitment zone is limited by the shape of *P*_*c*_(*i, j*): long-range effective interactions like in the diffusion scenario lead to more extended *P* (*M*) profiles compared to localized spreading as the NN case. In the 3D loop case, the *M* -state is able to stably propagate distally thanks to enriched 3D contacts with the painter area of sequence-specific recruitment.

**Figure 5.**
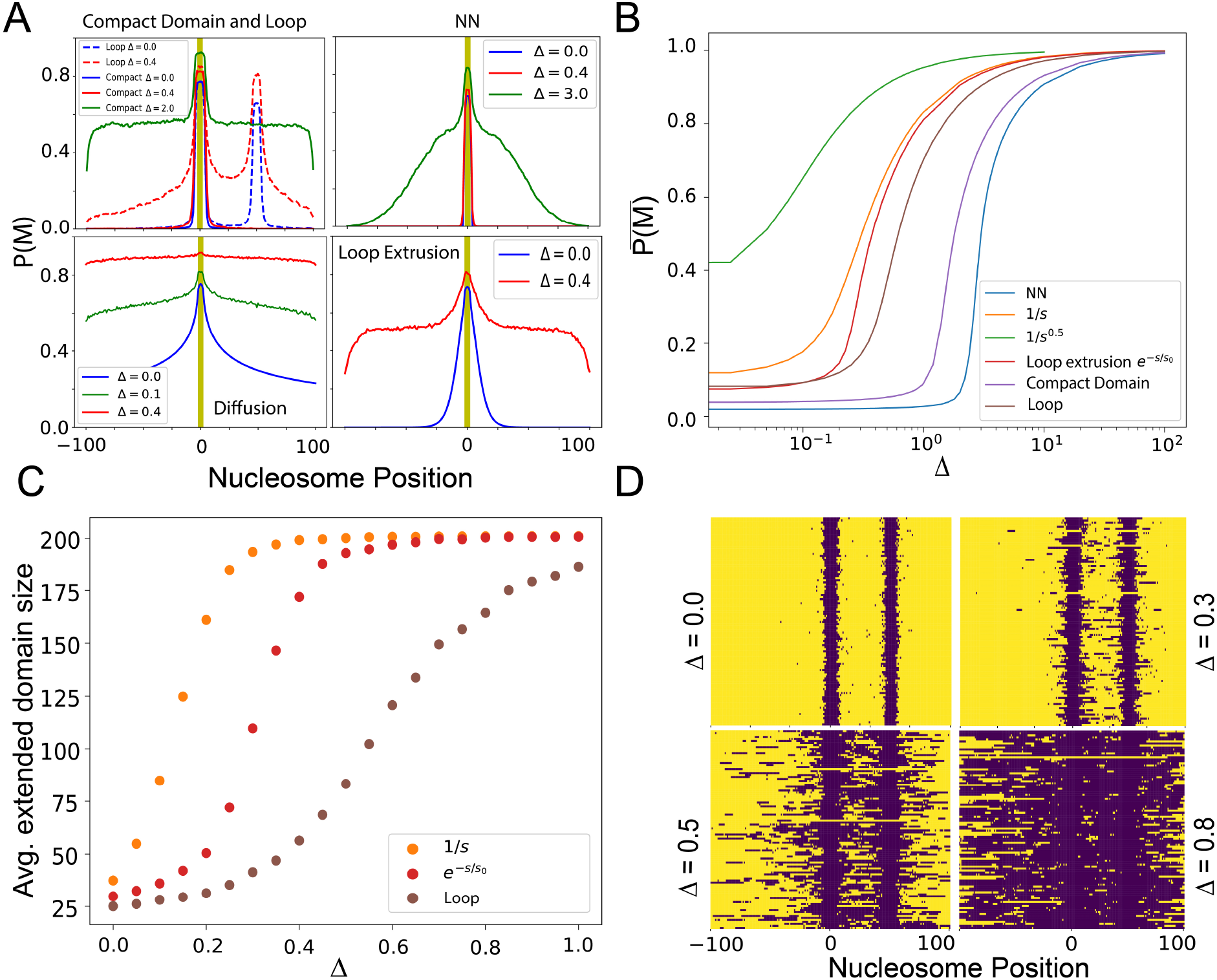
Spreading probability and percolation. (A) Virtual Chip-seq profiles of the M-state. *P* (*M*) as a function of the position around the 5 nucleosomes-long region where HMEs are recruited (yellow) for different *P*_*c*_(*i, j*) (see Materials and Methods) and for *k/k*_0_ = 1, *ε* = 0.6 with Δ = 0 (painter mode, Blue) and Δ *>* 0 (reader-writer mode, Red and Green). (B) Average value 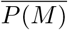 of *P* (*M*) inside the whole genomic region in the reader-writer mode as a function of Δ (*k/k*_0_ = 1, *ε* = 0.6) for different *P*_*c*_(*i, j*). (C) Average size of the most extended generalized domain as a function of Δ for three situations described in (B). (D) Illustration of the percolation of the system. For four different Δ values in the loop case (brown curves in (B) and (C)), most extended generalized domains observed in different configurations (100 examples) are stacked together to show cell to cell variability (all nucleosomes belonging to a domain are colored in blue).

In the reader-writer mode (Fig. 5A, Δ *>* 0), as observed in the previous section, state-specific recruitment allows a facilitated spreading and modifies the shape of *P* (*M*). For a given reader-writer strength Δ, scenarios with longer-range *P*_*c*_(*i, j*) are more impacted. However, for all scenarios, it exists a critical Δ value (Δ_*c*_) above which the *M* -state spreads over the whole region (Fig. 5B), more local *P*_*c*_(*i, j*) shapes (eg, NN or compact domain cases) exhibiting larger Δ_*c*_ values. Such transition driven by the reader-writer capacity of HMEs resembles actually to a generic percolation transition [27] where the system starts to be fully connected (or percolated) [72]. In our context, it corresponds to a situation where all the M-state nucleosomes form a unique expanded contact domain from which all the remaining U-state nucleosomes can be reached via *P*_*c*_(*i, j*) to allow spreading (see Materials and Methods, Supplementary Fig.5,Fig.10). To quantify this, for each configuration, we look how the most expanded M-state contact domain is connected to the unmodified regions. Fig. 5C shows the evolution of the average size of such extended generalized contact domains (containing U- and M-state nucleosomes) for three different scenarios. The global epigenomic state becomes percolated when this size approaches 200, meaning that the extended domain covers the whole genomic region (Fig. 5D). Interestingly, a full overall *M* -state 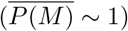 is not required to reach percolation, in particular for long-range spreading mechanism. For example, in the *P*_*c*_(*i, j*) = 1*/*|*j* − *i*| case, percolation transition occurs at Δ_*c*_ ≈ 0.2 where 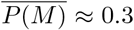 (Supplementary Fig. 10).

### 3.6 The reader-writer mode may lead to epigenetic memory

Having characterized how the various writing modes establish typical epigenomic profiles from the painter region, we now investigate how these different modes impact the “epigenetic memory” of these states. By definition, epigenetic memory stands for the ability of maintaining a given transcriptional or chromatin state in the absence of the initial, sequence-dependent stimulus (e.g. transcription factors or HMEs).

For such purpose, focusing on a 3D spreading mechanism (*P*_*c*_(*i, j*) = 1*/*|*j* − *i*|), we follow the time evolution of the mean genomic M-state 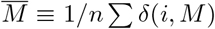 once the sequence-specific recruitment of painters have been released (for examples of individual trajectories, see Fig. 6C). In Fig. 6A, we plot the ensemble average-value 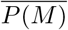 of 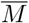 for different situations. In the simple painter mode (Δ = 0, green line), the M-state relaxes very quickly to the U state. Indeed, in absence of recruited writers in the painter area, only transitions from M to U are possible and the global state decays exponentially with a characteristic time 1*/k*_0_. Similar behaviors are observed in the booster-painter mode (Supplementary Fig. 11).

**Figure 6.**
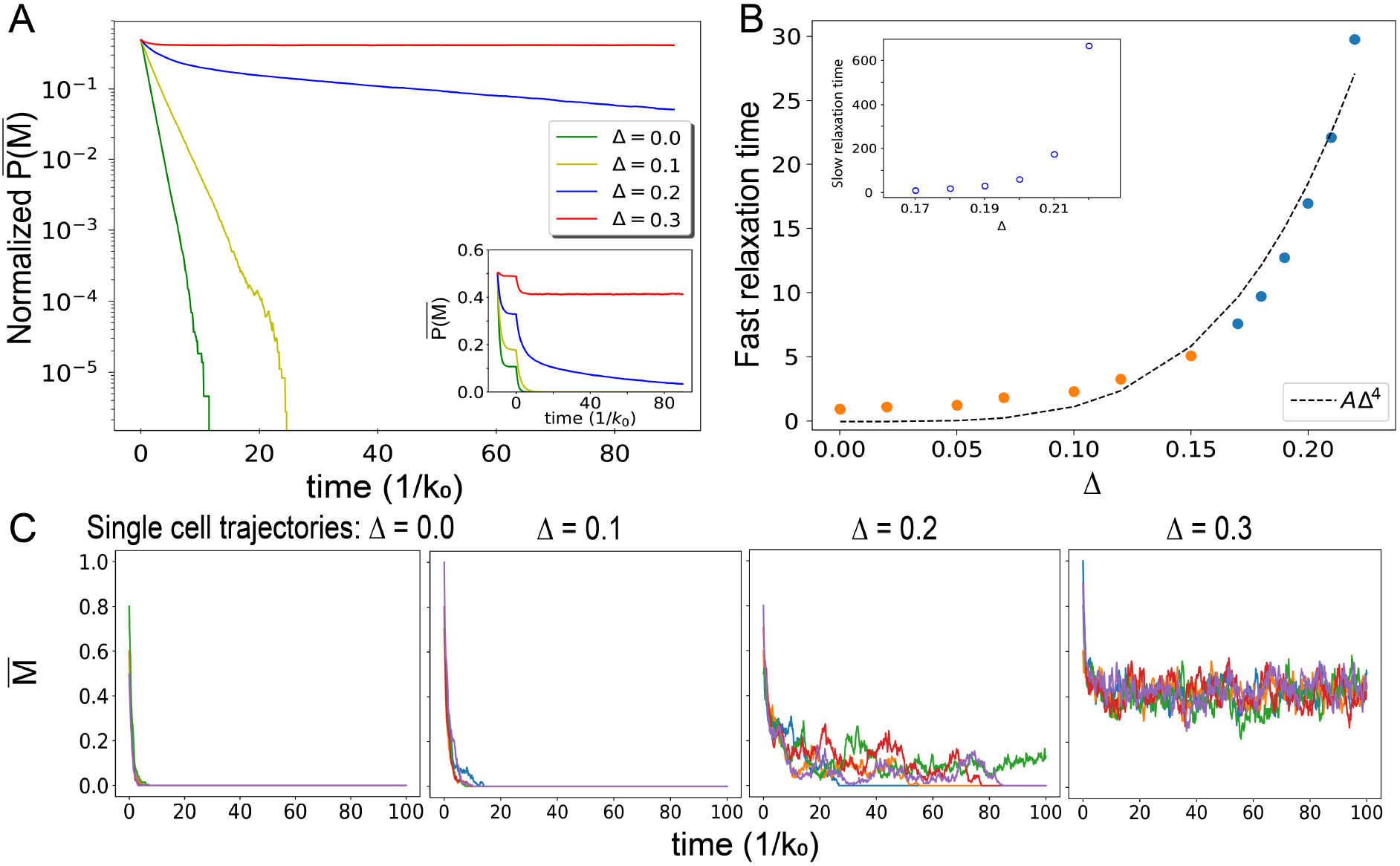
Epigenetic memory: (A) Normalized time-evolution of 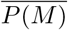 fter the unbinding of sequence recruited painters at *t* = 0 for different values of Δ and with *k/k*_0_ = 1, *ε* = 0.6. Inset shows the non-normalized evolution. (B) Fast (main) and slow (inset) relaxation times as a function of Δ obtained by fitting the curve in (A) by one or two time-scale exponential decay (see text). The fast time scales as ≈ Δ^4^. (C) Examples of “single cell” time-evolution of the M state decay (∑ *δ*_*i,M*_ */*201 vs *t*) for different values of Δ.

In the reader-writer mode, for low Δ values, the decay remains fast and exponential (Δ = 0.1 in Fig. 6A,C) with a relaxation time that increases with Δ (orange dots in Fig. 6B). Above a critical value, state-dependent recruitment is strong enough to stably maintain a M-state after HMEs unbind from the painter region (Δ = 0.3 in Fig. 6A,C) and to keep the memory of an initial (even small) M-state enrichment (Supplementary Fig. 12). The initial profile along the genome is lost and gives rise to an uniform spreading of the M-state (Supplementary Fig. 13).

Interestingly, for Δ-values just below this critical point, the relaxation dynamics is better described by a two time-scale exponential (Supplementary Fig. 14) kinetics (Δ = 0.2 in Fig. 6A,C): a fast initial decay following the trend of the low-Δ regime (blue dots in Fig. 6B) and a slow decay at larger time (inset in Fig. 6B). In this regime, random *M* → *U* conversions dominate but the state-dependent recruitment allows self-maintenance of a coherent M-state for long time periods before reaching a global, absorbing U-state 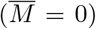 from which the system cannot escape as also observed by [27]. Such maintenance time-period is very stochastic (Δ = 0.2 in Fig. 6C) while, in the low and high Δ-regimes, convergence to steady-state behaves quite uniformly.

### 3.7 Maintenance of a confined chromatin state requires chromatin compaction and enzymatic titration

In the previous section, we showed that epigenetic memory was possible for strong-enough state-specific recruitment. However, in this regime of parameter, the spreading of the M-state along the genome is not constrained [39], leading to unconfined memory [24] (Fig. 7A). Of course, experimentally, epigenomic domains are confined to specific regions in the genome and do not spread ubiquitously [10, 16]. In this section, we investigate what could be the minimal changes to our model to support confined memory and the maintenance of a localized M-state even in absence of sequence-specific recruitment.

**Figure 7.**
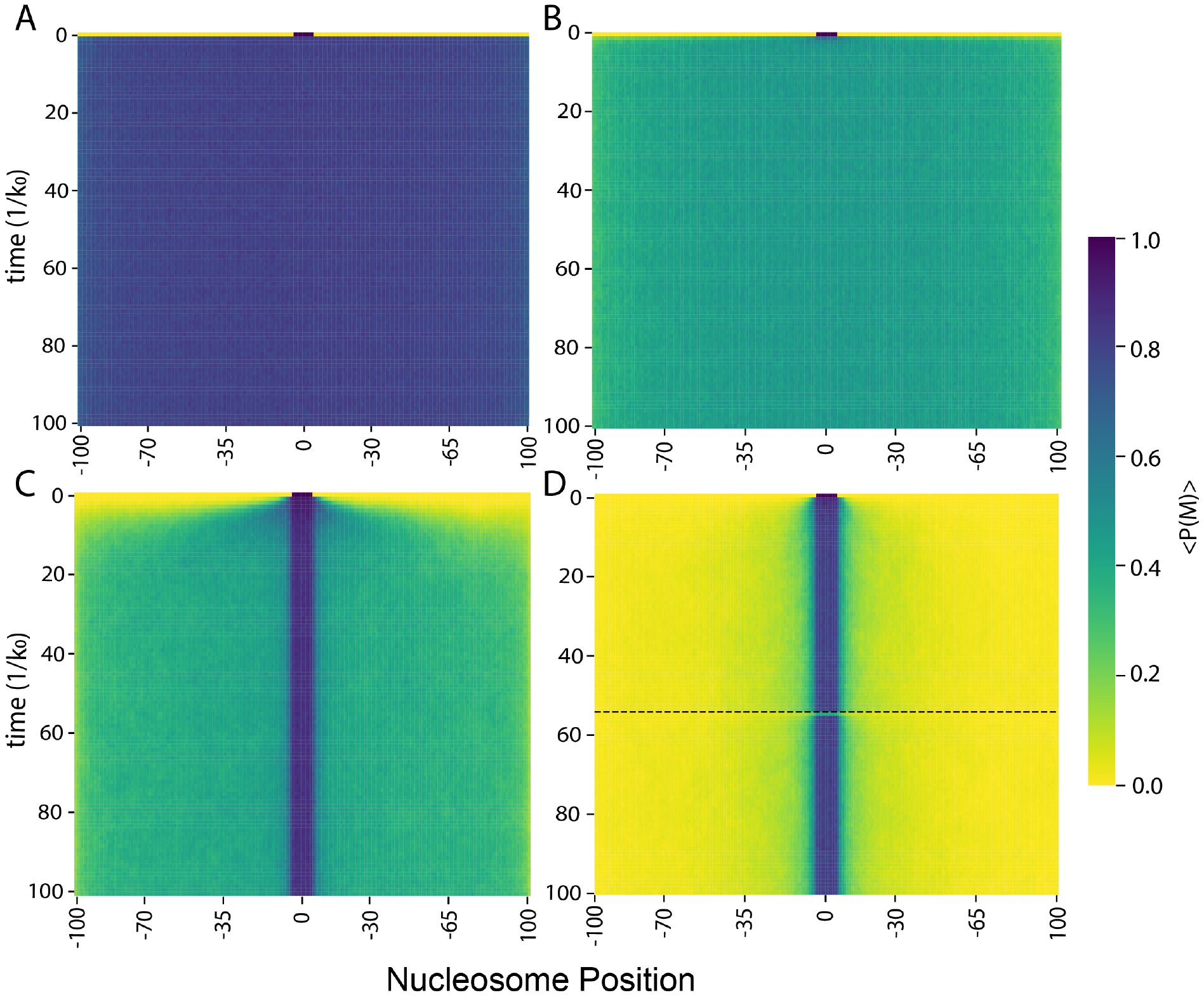
Confined epigenetic memory. (A) Evolution of *P* (*M*) in the enzyme non-limiting regime *N* ^*tot*^ = 45. The initial configuration is a domain of modified states around nucleosome position 0 and there is no sequence-specific recruitment for *t >* 0. *k/k*_0_ = 2, *ϵ* = 0.9. (B) As in (A) but with enzyme limitation, 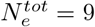. (C) As in (A) but with 3D compaction of the domain (Supplementary Fig.9A shows the corresponding contact probability matrix), in large 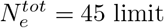. (D) As in (A) but with enzyme limitation 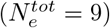 and 3D compaction. Modified state domain remains confined even under replicative dilution (*T*_*cyc*_ = 55*/k*_0_, dashed black line).

Our simple epigenomic model assume implicitly that the number of HMEs is not limiting in the system. Recent quantitative proteomics experiments however suggest that such assumption might not be satisfied for some epigenomic marks like the Polycomb/H3K27me3 system during fly embryogenesis [85]. Therefore, a mechanism that could possibly restrict the spreading of M-state might be enzyme limitation. In the model, under the approximation of fast binding-unbinding enzyme kinetics at M-state nucleosomes, it is possible to account for enzyme titration via an effective Δ parameter that depends explicitly on the total number of HMEs 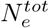 and on the current fraction of bound HMEs (see Materials and Methods), high fractions being associated with low effective state-dependent recruitment. Limiting the number of HMEs thus reduces the spreading efficiency (Supplementary Fig. 4), however in ranges of parameters that support epigenetic memory, the long-range-spreading activity of bound HMEs still lead to unconfined epigenomic domains (Fig. 7B).

As long-range spreading via the trans-activity of HMEs seems to promote unconfined memory, another possible mechanism to restrict spreading might thus be to modulate the 3D communication between loci [37]. Some epigenomic marks are associated with architectural proteins that may indeed impact chromatin organization. For example, PRC1 and HP1 that can bind H3K27me3- and H3K9me2/3-marked chromatin respectively, are known to promote chromatin compaction in vitro [86, 87] and in vivo [88–90]. To explore how 3D compaction of domains would influence epigenetic memory, we introduce a nucleosome-nucleosome contact probability *P*_*c*_(*i, j*) that is consistent with the formation of a compact domain around the painter region (see Materials and Methods) with *P*_*c*_ inside the domain being stronger than outside (Supplementary figure 9). In the limit of high 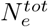 (Fig. 7C), such 3D organization clearly facilitates the maintenance of a stable M-state inside the compacted region even in absence of sequence dependent stimulus, but exhibits “flooding” of the M-state outside this region with time due to the residual spreading between nearest-neighbor sites. However, when we couple 3D compaction and enzyme limitation, the initial domain remains very stable with limited flooding even under strong perturbation like replication (Fig. 7D), suggesting that both ingredients can lead to confined epigenetic memory.

### 3.8 Chromatin state dynamics regulates transcriptional noise

Histone modifications are likely to play a role in the regulation of genome accessibility thereby impacting transcriptional activity of genes [3]. To understand how gene expression may be affected by the dynamics of chromatin states, we consider a simple toy model where the transcriptional state of a gene depends on the current mean M-state proportion 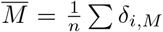 over a 1kbp-wide (*n* = 5) region that would represent the gene promoter. We consider in this section that the state M is a repressive state (e.g. constitutive heterochromatin via H3K9me2/3) and that the transcription rate is inhibited by the presence of modified nucleosomes at the promoter 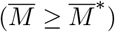 in a switch-like manner [22](Fig. 8A, see Materials and Methods).

**Figure 8.**
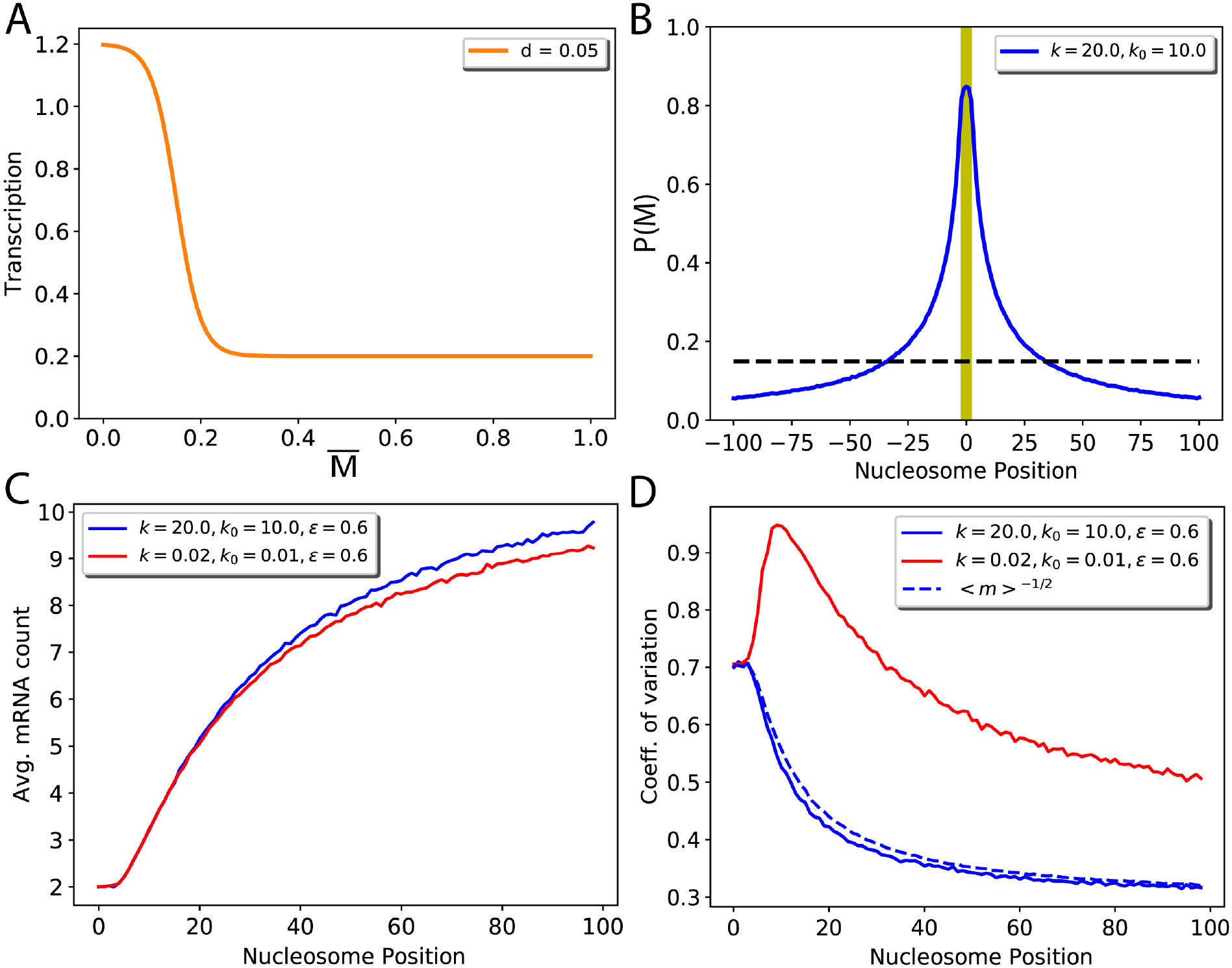
Chromatin state dynamics and transcription: (A) Cooperative switch model: Transcription rate as a function of M state proportion 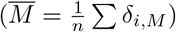 with steepness *d* = 0.05. (B) *P* (*M*) profile for *k/k*_0_ = 2, *ε* = 0.6. Black dotted line indicates the value of *P* (*M*) at the transcriptional switch (*P* (*M*) = *M*^∗^,see (A)). (C) Average mRNA count for the two different epigenomic dynamics as a function of the distance of the gene from the painter region. (D) The coefficient of variation of the mRNA count for the same parameters as in (C). The dashed line shows 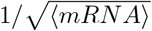 with ⟨*mRNA*⟩ taken from panel (C).

To only focus on the epigenetics-transcription relation, we consider a simple painter mode (as in Fig. 2) with *k/k*_0_ = 2 and investigate the transcriptional properties of a gene as a function of the proximity of its promoter to the painter region (Fig. 8). We observe that the steady-state average number of mRNA per cell ⟨*mRNA*⟩ for this gene is larger for more distant promoters (Fig. 8C), qualitatively mirroring the decrease in *P* (*M*) (Fig. 8B). Interestingly, for the same *relative k/k*_0_ value (and thus the same average M-state profile, see Eq. 3, Fig. 8B), the average transcription level depends on the *absolute* values of *k* and *k*_0_ that control the kinetics of M state dynamics, comparatively to the transcription dynamics that is itself driven by the transcription and mRNA degradation rates (see Materials and Methods). We detect that slow epigenetic dynamics (red line in Fig. 8C) tend to favor a slightly more repressed state compared to fast dynamics (blue line in Fig. 8C). Such kinetic effect is more striking when considering the intrinsic stochastic fluctuations of gene expression by estimating the coefficient of variation (CV) (Fig. 8D), defined as the ratio between the standard deviation of the corresponding steady-state distribution of mRNA number per cell and its average value, a high CV meaning a noisy, highly fluctuating gene. For fast dynamics (blue line in Fig. 8D), relative fluctuations decrease with the distance from the painter region. CV follows 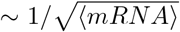 (dashed blue line in Fig. 8D), characteristic of the intrinsic transcriptional noise found in elementary gene expression models [93]: lowly expressed gene being relatively more noisy. For slow dynamics (red line in Fig. 8D), we instead observe a sharp increase up to a distance *s*^∗^ where fluctuations become maximal, followed by a gradual decrease at larger distances; fluctuations remaining always larger than in the fast dynamics case. Indeed, in our transcription rate model (Fig. 8A), close to the transition point 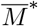, small variations of 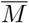 result in large deviations for the transcription rate. If the epigenetic dynamics is faster than mRNA production and degradation rates, the transcription level cannot adjust to the rapid fluctuations of 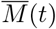 that are filtered by the slow mRNA dynamics [94]. However, if the chromatin dynamics is much slower, mRNA level adapts to the current, slowly fluctuating M-state. Hence expression will stochastically switch between a repressed 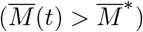 and expressed state 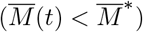 leading to large fluctuations of expression. The peak observed in CV in this case thus translates the interplay between such ultra-sensitivity to epigenetic fluctuations with the more standard dependence of the noise to ⟨*mRNA*⟩ (fast chromatin dynamics case). Biologically, this strong propagation of epigenetic fluctations to the gene expression level may be related to the well known variegation phenomenon where genes inserted near heterochromatin domains may have variable transcriptional pattern, depending on the distance of their insertion sites to the heterochromatin domains [95].

### 3.9 Applications to diverse biological contexts

Above we studied systematically the behavior of our generic model of epigenomic regulation and its consequences on epigenetic memory and transcription. In the next, we will describe three applications of this framework to specific biological situations.

#### 3.9.1 Phosphorylation of *H*2*AX* around double-stranded breaks

During our analysis of the simple painter mode (Fig. 2), we showed that there is a direct relationship between the distribution of marks around the painter binding sites and the local 3D chromatin organization. This 3D↔1D relationship has been evidenced experimentally in the context of DNA double-stranded breaks (DSB) in humans [70]. After a break had occured, variant histones H2AX present in the chromatin are phosphorylated (the so-called *γ*H2AX mark) by the ATM kinase recruited at the DSB site. The correlative analysis of the 4C and *γ*H2AX Chip-seq signals around DSBs clearly reveal high similarities (e.g. see Fig. 1A in [70]), suggesting that the spreading by painter mode is at work in that system. In Fig. 9A, we extracted the average *γ*H2AX profile observed around DSBs (red dots) from [70] that extends over Mbps. Using the analytical formula for a simple painter model (Eq. 3) with a painter region localized at the DSB and *P*_*c*_(*s*) ∼ 1*/s* the typical average contact probability found in human cells [61], we were able to fit very well the experimental data (blue line in Fig. 9A, L2-distance = 1.4 ∗ 10^*−*4^)(see Materials and Methods). This leads to one identifiable parameter (*k/k*_0_)*ε* = 1213. In the original article, Arnould *et al*. suggested that the loop extrusion mechanism might be directly implicated in the spreading mechanism [70]. To test this alternative hypothesis, we perform a similar inference using the loop extrusion-like spreading probability (*P*_*c*_(*s*) ∼ exp[−*s/s*_0_]) which leads to a less precise fit even if the model has an additional parameter (processivity *s*_0_ ∼ 150 − 200kbp) (green line in Fig. 9A, L2-distance= 5.1 ∗ 10^*−*4^). Note that the diffusion-like spreading mechanism (*P*_*c*_(*s*) ∼ 1*/s*^0.5^) is also inconsistent with the experimental data (orange line in Fig. 9A, L2-distance= 9.0 ∗ 10^*−*4^). These results differ from a similar analysis performed by Li et al. [91] suggesting, using a two-state mathematical model neglecting turnover rate of the mark, that Tel1, the yeast homolog of ATM, spreads phosphorylation around DSB via a loop extrusion-like mechanism while within our framework the best prediction (minimum L2-distance) is given by the spreading-by-3D-contact mechanism (*P*_*c*_(*i, j*) = 1*/*|*j* − *i*|)).

**Figure 9.**
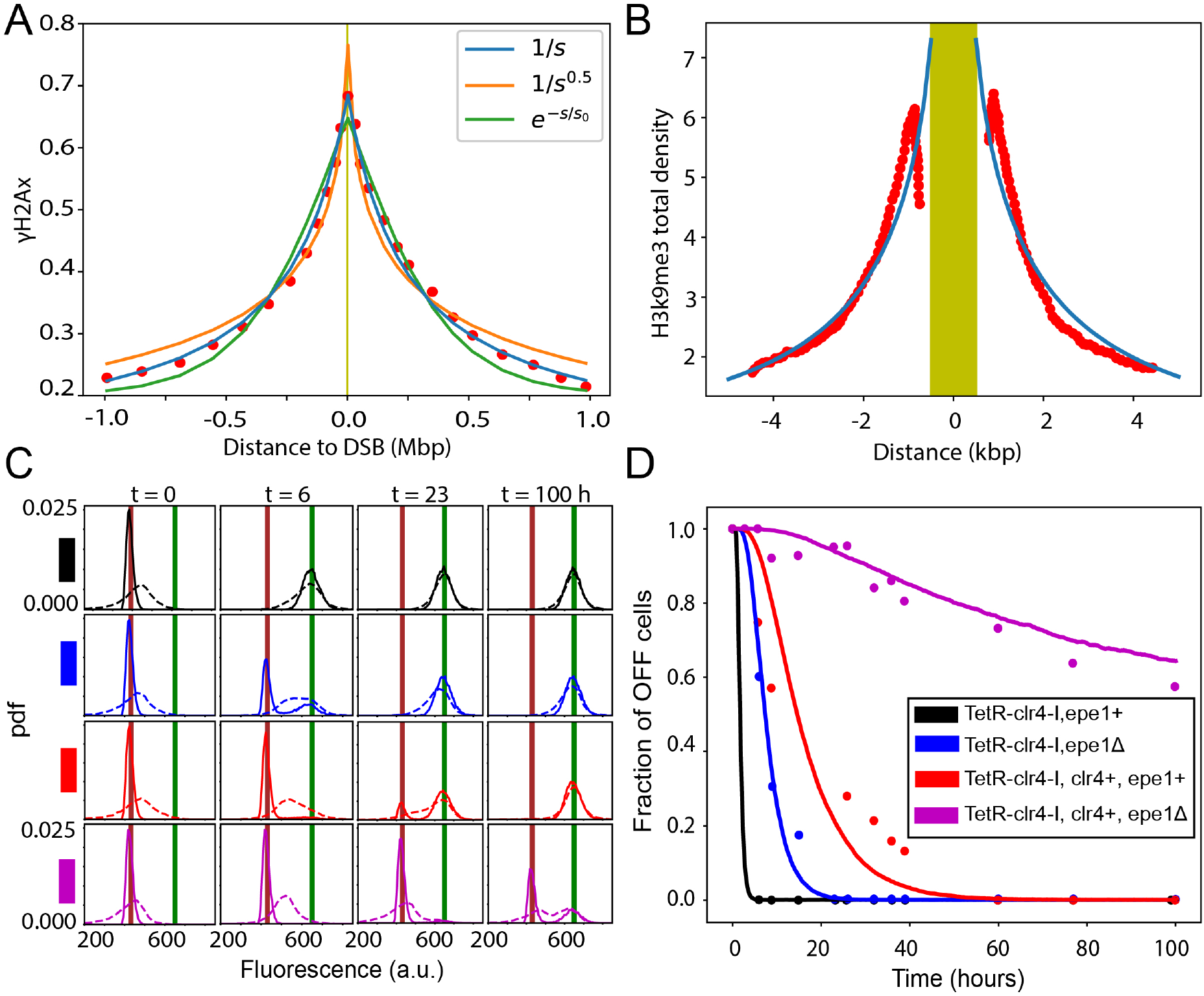
(A) Experimental phosphorylation profile (dots) around a DNA double strand break [70] and the corresponding model prediction (i) *S*(*i*) = 0.58*P* (*M*_*i*_) + 0.11, (*k/k*_0_)*ε* = 1213 *P*_*c*_(*i, j*) = 1*/*|*i* − *j*| (blue) (ii) *S*(*i*) = 0.85*P* (*M*_*i*_), (*k/k*_0_)*ε* = 29.6, *P*_*c*_(*i, j*) = 1*/*|*i* − *j*|^0.5^ (orange) (iii) *S*(*i*) = 0.9*P* (*M*_*i*_) + 0.2, (*k/k*_0_)*ε* = 1, 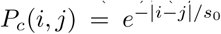, *s*_0_ = 1050 (green). (B) Experimental H3K9me3 density (dots) flanking a transposable element in mouse ES cells [74] and the corresponding model prediction *S*(*i*) = 10.75*P* (*M*_*i*_) + 0.1 (full line). (C) Heterochromatin maintenance in fission yeast. Distributions of fluorescence expressed from a reporter gene at specific time points after painter release at *t* = 0 for different strains (experiments: dashed lines [10], model predictions: full lines). Brown and green vertical lines mark the mean fluorescences in the OFF and ON states respectively. (D) Experimentally-measured (dots) and predicted (full lines) time-evolution of the proportion of OFF cells in the population for the various strains.

#### 3.9.2 Heterochromatin formation around retrotransposons

An entirely different system where the simple painter mode may also be operative is the spreading of H3k9me2/3 modifications over few kbps around retrotransposons in mouse ES cells by Setdb1 [74, 96], a lysine methyltransferase recruited by the KRAB–Zinc Finger Protein KAP1 [97]. As in the previous example, we can fit the experimental Chip-seq profiles (red dots in Fig. 9B) by a simple model with a painter zone corresponding to the typical size range of a transposable element ∼ 1*kbp* [75] (blue line in Fig. 9B). In this case, we found (*k/k*_0_)*ε* = 0.78. Note that these predictions were done without invoking the reader-writer mechanism which might be relevant in other heterochromatin contexts (see below).

Compared to *γ*H2AX, the spreading of H3K9me2/3 is much less efficient leading to smaller domains (kbp-vs Mbp-wide). This may translate fundamental differences in the spreading and turnover rates as histone methylation maintenance in mammalian cell lines could be slow (∼ 10-20 hours) [29, 59] while establishment of *γ*H2AX foci is much faster (∼ 1-30 min) [98, 99].

#### 3.9.3 Memory of heterochromatin in fission yeast

Finally, we apply our chromatin state dynamical model to quantitatively characterize the stability of epigenetic memory for fission yeast heterochromatin. In *Schizosaccharomyces pombe*, establishment of H3K9me2/3 domains is in part regulated by the balance between the methyltransferase activity of Clr4, its state-dependent recruitment by H3K9me histones and the demethylase-like action of Epe1 [10, 100]. In [10], a heterochromatin domain is established via the sequence-specific recruitment of Clr4 at an ectopic locus containing a fluorescent reporter gene, leading to the repression of the gene. Epigenetic memory of this domain and involved mechanisms are then characterized by releasing sequence tethering and by tracking the progressive re-activation of the gene as a function of time (full lines in Fig. 9C). In particular, they considered four different strains: (i) *TetR* − *Clr*4 − *I, epe*1+ with a mutated Clr4 (*Clr*4 − *I*) with no reader-writer property and a normal demethylase-like activity (*epe*1+), the model analogue being Δ = 0, *k*_0_ = *x*; (ii) *TetR* − *Clr*4 − *I, epe*1Δ same as (i) but without demethylase-like activity (*epe*1Δ) (model equivalent - Δ = 0, *k*_0_ *< x*); (iii) *TetR* − *Clr*4 − *I, Clr*4+, *epe*1+ with wt-like Clr4 and Epe1 activities (Δ *>* 0, *k*_0_ = *y*); and (iv) *TetR* − *Clr*4 − *I, Clr*4+, *epe*1Δ as in (iii) but in *epe*1Δ background (Δ *>* 0, *k*_0_ *< y*). They observed that memory of a repressed state was enhanced by the reader-writer module of Clr4 and the absence of Epe1 (dots in Fig. 9D).

To rationalize these experiments with our quantitative framework, we simulate the experimental memory assay like in Fig. 6 but for various values of Δ and *k*_0_ and by tracking the transcriptional activity of the region using the model described in Fig. 8 (see Materials and Methods). More precisely, we follow the experimental protocol and monitor the time-evolution of the distribution of fluorescence of the reporter gene within the cell population (Fig. 9C), from which we can define the proportion of cells that are still repressed (“OFF”) by heterochromatin (see Materials and Methods) (Fig. 9D). The predicted time-evolution of this proportion for each scanned (Δ,*k*_0_) value is then quantitatively compared to the experimental data obtained for each of the four strains (Supplementary Fig. 6, see Materials and Methods). For strain (i), gene activation is fast (Supplementary Fig. 15, black dots in Fig. 9D) and data are compatible with a simple painter mode with high turnover rate (*k*_0_ *>* 0.007 *min*^*−*1^, Δ = 0) (Supplementary Fig. 6A, and black line in Fig. 9D), consistent with the presence of Epe1 and the absence of reader-writer recruitment which, we demonstrated in Fig. 6, is essential for epigenetic memory. For strain (ii), the gene de-repression is slower than in (i) (blue dots in Fig. 9D) but can still be well captured by a simple painter mode (Δ = 0) (blue line in Fig. 9D) associated with decrease of *k*_0_ by at least 6 fold (Supplementary Fig. 6B), consistent with absence of Epe1 in this strain and thus with less turnover. In both backgrounds (iii)(*epe*1+) and (iv)(*epe*1Δ), the reader-writer capacity of Clr4 strongly enhances memory and slows down gene activation (red and purple dots in Fig. 9D). This behavior can only be reproduced (red and purple lines in Fig. 9D) by a reader-writer mode with Δ = 0.004 (Supplementary Fig. 6C,D). Interestingly, for strain (iv) where memory is maximal, the model predicts that the biological system is in a parameter range just below the critical regime of the reader-writer mode (like the blue line in Fig. 6A). In this regime, the epigenetic state is long-lastly maintained until rapid removal of the marks (Δ = 0.2 in Fig. 6C), leading to a stochastic switch in transcriptional activity and bimodal distributions for protein levels (Fig. 9C, purple at t = 100h), as observed experimentally [10].

## 4 DISCUSSION AND CONCLUSION

In this work, we developed a unified mathematical two-state model of epigenomic regulation, integrating key mechanisms like reader-writer processes. Our simple framework states that the establishment and maintenance of an epigenetic state result from the recruitment and spreading activity of histone modifying enzymes (HMEs). Recruitment of HMEs can be mediated by specific genomic sequences or by the local epigenomic state. Spreading encompasses an on-site action of the HMEs and a long-range, trans activity modulated by the local chromatin state.

In particular, we systematically studied three generic modes of regulation that recapitulate most of the experimentally-known epigenetic systems: (i) a simple painter mode (Fig. 2) in which HMEs are targeted to specific genomic regions and spread via their trans-activity around these binding sites; this mode may be representative of the regulation of small, local epigenomic domains like acetylation marks (e.g. H3K27ac or H3K9ac) around promoters and enhancers; (ii) a boosted-painter mode (Fig. 3) in which recruited HMEs have an enhanced activity if bound to specifically-modified regions; this mode may account for the allosteric boost observed for PRC2 in presence of H3K27me3 [17]; (iii) a reader-writer mode (Fig. 4) in which HMEs may also be recruited by specific epigenetic signals; this mode may capture the regulation of extended chromatin domains like heterochromatic regions (e.g. H3K9me2/3).

In the simple and boosted-painter modes, we found that there is a direct, simple relationship between the binding profiles of HMEs, the chromatin organization and the profile of epigenomic state (Eq. 3). Verification of this relation using experimental measurements of these three information (e.g. bulk Chip-seq and Hi-C data) in wild-type-like conditions may be a strong evidence for one of these two spreading modes [29]. A perfect illustration of this is our application to *γ*H2AX around DSBs in which the experimental profile is well fitted by the model (Fig. 9A) suggesting a simple or boosted-painter mode with a spreading-by-3D-contact scenario rather than a more complex cohesin-mediated mechanism [70].

Distinguishing the boosted-from the simple-painter mode would then require additional information like for example to estimate the spatial correlations between chromatin state that have very different signatures between the two modes (Fig. 2C *vs* 3C). However, this information is currently very difficult to access experimentally as it would require a single-cell assay to estimate covariations of chromatin states; but recent progresses in single-cell Chip-seq experiments [105, 106] may open new venues. Another promising application of Eq. 3 in the simple/boosted-painter modes is that it can be employed to address the inverse problem of inferring the HME binding sites knowing the 3D chromosome organization (e.g. via Hi-C) and the Chip-seq profile of epigenetic states, or of estimating the chromatin folding properties from HME and histone mark Chip-seq profiles. Actually, the latter strategy was recently used by Redolfi *et al*. [82], in the so-called Dam-C technique, to extract 4C-like contact probability information based on the spreading of Dam-mediated DNA methylation from a painter region.

In the booster-painter and reader-writer modes that both involve a “reader” capacity of HME, we observed phase-transitions and critical behaviors via respectively the strengths of boosting activity (*r*) and of state-dependent recruitment (Δ) (Figs. 3B, 4B). These behaviors are driven by the effective positive feedback loops and cooperative effects [18] that emerge from the enhanced spreading efficiency of some HMEs (more activity or more recruitment) if the histone modification they catalyze is already present locally. These effects are more important in the reader-writer mode and lead to qualitatively different epigenomic profiles, with extended domains around the binding peaks of HME (Fig. 4A), and distal spatial correlations (Fig. 4C). In particular, we found that these profiles are very sensitive to the underlying spreading capacity of HMEs [101], longer-range mechanisms facilitating the percolation of the system into large contact domains at low state-dependent recruitment (Fig. 5C) [27]. Eventually, these domains may be maintained in the absence of genomic bookmarking (Fig. 6A) for strong enough state-dependent recruitment. However, our analysis of heterochromatin memory in fission yeast (Fig. 9C,D) suggests that such self-sustainable memory may not be occurring for H3K9me2/3 chromatin domains in wild-type conditions and that (weak) nucleation, sequence-dependent signaling might still be needed for maintaining a stable epigenetic landscape [37, 39]; even if a reduction in the effective histone turnover rate (e.g. by a lower demethylase activity as in the *epe*1Δ mutants in [10] or by longer cell-cycles [22, 30]) might trigger the system near to the critical point and may lead to long-term memory [27], but potentially also to high sensitivity to external cues [32]. As shown by previous modeling works [24, 27], we also confirmed that purely local spreading mechanisms (as the NN scenario) would require too strong feedback and state-dependent recruitment to stabilize extended, coherent epigenomic domains during a long time period, even if being able to maintain them over few cell cycles [103].

One theoretical issue however with standard reader-writer models with long-range spreading is the difficulty to stabilize finite-size epigenomic domain without genomic bookmarking and to avoid unconfined spreading at long-time scale, as already pointed out by Erdel and Greene [24] and Dodd and Sneppen [19]. While strong insulators or barriers preventing local spreading [19, 37] or/and the formation of compact 3D domains [37] may slow down but not prevent uncontrollable spreading, we showed here that the combination of 3D compartmentalization with enzyme titration may lead to ultra-stable confined memory of small domains even under strong perturbations like replication (Fig. 7D), as also previously observed by Sandholtz *et al*. using an explicit polymer model of epigenetic regulation with controlled HMEs concentrations [102]. This may be particularly relevant for the Polycomb regulatory system as it involves HMEs (PRC2) in limited numbers, as measured recently during fly embryogenesis [85] and evidenced also in mammals [107], and is associated with PRC1, a protein complex promoting the compaction of H3K27me3-tagged regions [31, 86]. Similarly, maintenance of confined silenced domains in yeast may rely on the combination of their spatial compartmentalisation via Sir3-mediated self-attraction [108] and Sir4-driven tethering to the nuclear envelop [109]) and of the titration of the Sir2 deacetylase [110, 111]. Note that, in our model, 3D compartmentalization was accounted via an attractive interaction between M states, mimicking PRC1, HP1 or Sir3 modus operandi. However, additional architectural mechanisms such as the spatial co-association of boundary elements are likely to be at work: in mammals, the stability of Polycomb domains may be reinforced by contacts between CTCF insulator sites [112] possibly via the cohesin-SA2-mediated loop-extrusion process [113]; in yeasts, colocalization of TFIIIC-binding boundary elements participate to chromatin domain integrity [43, 48]. Additionally, the presence of antagonistic chromatin states domains may act as competitive barriers to epigenomic spreading [6] but also may favor 3D compartmentalisation by strengthening phase separation [114]. Actually, all kind of processes that reinforce *intra-* vs *inter-* contact domains is expected to promote confined memory by limiting long-range “contamination”: in a condition of limited number of HMEs, increasing compartmentalization will further increase their local concentration (by sequestration), leading to a stronger stability and weaker pervasive long-range spreading (Fig. 7D), in agreement with recent experimental observations in fission yeast suggesting that a “critical density” of H3K9me3 is required for the stable inheritance of confined heterochromatin [104].

To go beyond the mere description of epigenomic regulation, we also proposed a simple mathematical model describing the impact of chromatin state dynamics on gene expression [22]. Using this generic and modular framework, we quantified the role of epigenetic fluctuations in transcriptional noise. It allowed us to investigate positional variegation as well as epigenomic memory in terms of gene expression in addition to histone marks. In particular, this makes possible to analyze and interpret the transcriptional outputs of epigenetic-related experiments. For example, using this framework, we quantitatively described heterochromatin assays by fitting gene reporter expression distribution versus time and strains. Introducing a feedback of transcription on the epigenomic dynamics via the recruitment of HMEs [26, 28] or changes in turnover rates [115] may allow a finer description of the interplay between epigenomics and transcription.

Previous mathematical models have also addressed various aspects of the establishment and maintenance of epigenomic information. Several works [18, 20, 26, 32, 118] have focused on bivalent regions, regulated by reader-writer and long-range spreading mechanisms, that can alternate between different bistable, coherent states (active or inactive). While our framework can be easily generalized to account for such multistability, the scope of our study was to specifically investigate the interplay between reader-writer processes, spreading capacity and nucleation mechanisms. In that context, our approach generalizes or complements previous two-state descriptions of epigenome regulation, like the 3D looping models of Erdel and Greene [24] and Katava et al. [39] and the stochastic model by Ancona et al. [27]. 3D looping models with sequence-dependent recruitment and a long-range spreading mechanism based on 3D contacts are by essence very similar to our simple painter and reader-writer modes. In [24], authors discussed the role of looping (vs NN scenario) in epigenomic profiles and of reader-writer feedback in memory and proposed that non-homogeneous state-specific recruitment may lead to stable confined domains. In [39], they investigated the interplay between spreading dynamics and chromatin motion, highlighting the role of nucleation in the maintenance of stable domains in a regime of fast epigenetic spreading. In their stochastic model, Ancona et al. [27] studied a reader-writer-mode like model in absence of nucleation and showed that local mark turnover coupled to long-range spreading mechanisms may lead to percolation transition and bistability.

In our work, we focused on a single painter site, but to go further the painter model may provide a very interesting framework to specifically address the pivotal role of the recruitment painter sites in globally shaping the epigenome. In particular to investigate how their sizes, strengths or genomic distributions influence epigenomic regulation [101]. Interestingly, within some epigenomic domains, some sequences can recruit HMEs while being unable to nucleate chromatin states on their own [7]. These elements may actually act as secondary recruitment sites after the initial nucleation events at the strong, autonomous sites [116] and, in the presence of numerous weak sites, a stable domain can be formed even in the absence of strong sites [117]. Based on our results, an appealing hypothesis would be that, the numerous dispersed sites along the genomic domain might compensate their “individual” weakness by maintaining “collectively and cooperatively” a high local concentration of HMEs via long-range 3D “communication”.

One strong hypothesis that we made in order to focus on the painter and reader-writer processes and their consequences on epigenetic memory, was to neglect the interplay between several chromatin modifications [118, 119]. Indeed, the regulation of one silencing epigenomic mark often results from the competition with antagonistic activation marks. Such tug-of-war is driven by reader-writer processes complemented with reader-eraser mechanisms that actively remove opposite marks [118, 120]. This may lead to bistability between active and inactive chromatin states and participate in reinforcing epigenetic memory [18, 26, 121]. As for writers, recruitment of eraser enzymes may be sequence- or state-specific. For example, the UTX/JMJD3 demethylase is recruited by Trithorax-group proteins at active developmental genes to remove the repressive Polycomb H3K27me3 mark [14, 122]. In addition, the regulation of histone turnover may also participate in the erasing mechanisms. Indeed, genomic regions tagged with inactive modifications like H3K9me2/3 are know to display low histone turnover rate, while transcribed regions tends to exhibit higher rates [54, 115, 123]. All this may be responsible of complementary feedback loops in the epigenomic regulatory dynamics that finely tune the different levels of epigenomic modifications. For example, transcription may mediate such a loop not only by depending on the current epigenomic state, as we tentatively modeled in Fig. 8, but also, in return, by impacting epigenomic stability via an increase in histone turnover rate [26, 28].

Another simplification made in our work was to assume that the 3D chromatin organization (that mediates the long-range spreading) was fast compared to epigenomic-associated rates and was independent of the current chromatin state. While the former hypothesis may be satisfied for short genomic distances, for large, Mbp-wide, chromatin domains like pericentromeric, H3K9me2/3-tagged regions in higher eucaryotes, the dynamics of contact may be slow [58,124] leading to less efficient spreading [37,60]. As discussed above, heterochromatin domains (e.g. H3K9me3, H3K27me3) are often associated with architectural proteins (e.g. HP1, PRC1) that may impact on their compaction. Therefore, it introduces again a dynamical feedback loop between long-range epigenomic spreading and 3D chromatin organization: nucleation of a chromatin state drives its compaction that in turn facilitate spreading and maintenance [35, 87]. Recently, we and other groups investigated more carefully this coupling by developing a model that explicitly account for both 3D and 1D dynamics of chromatin [37–39, 102], highlighting the key role played by genome folding on epigenomic regulation. Combining the formalism developed here with such more detailed framework would allow to better characterize the structure-function relationship of chromatin including the formation and maintenance of confined chromatin domain.

To conclude the formalism developed here is generic and modular, as it provides a simple description of epigenomic regulation in terms of HMEs recruitment and enzymatic activity. It can be easily upgraded to include more chromatin modifications and cross-talks and feedback loops and can thus be contextualized to a variety of specific chromatin regulatory systems [29].

A very promising application of our painter model might be to explore how epigenomic (and corresponding transcriptional) dynamics might influence genome evolutionary processes. Of particular interest is the challenging question of the role of epigenome in the control of the integration and selection of transposable elements (TEs). TEs are mobile genomic elements that can be inserted at new genomic positions or can modify their location either via a “copy and paste” or in a “cut and paste” mechanism [75]. They comprise as much as 40 − 60% of mammalian genomes. For active retrotransposons, a specific class of TEs, host cells have developed defense strategies in order to limit their invasion and expansion and to maintain genome integrity. Most of these elements are targeted by epigenomic silencing mechanisms (e.g. H3K9me2/3) [125–127] that limit their expression and thus, in *fine*, restrict their capacity of “parasitic” transposition [128]. This silencing is usually achieved by site specific recruitment of HMEs such as Setdb1 and Su(var)3-9s that nucleate and further spread the H3K9me2/3 marks [125,129,130] within TE elements and beyond into the flanking region, as shown in Fig. 9B with the simple painter mode in mESC. Very similar spreading patterns have also been observed around several other TE families in two Drosophila species, *D. melanogaster* and *D. simulans* [131]. As expected and confirmed by our theoretical approach that couples epigenomic and transcriptional dynamics (Fig. 8C), such long-range spreading of repressive chromatin state from TE elements has been shown to mediate long-range silencing of flanking genes [132]. This is actually reminiscent of the well known phenomenon of position effect variegation (PEV) observed in various organisms, where the incidental repositioning of a normally expressed/repressed genes, by translocation or transposition, next to a hetero/euchromatin fuzzy domain boundary induces its stochastic repression/expression depending on the genomic distance between the insertion and the boundary [95, 133, 134]. On one hand, there is a generic selection against TE transposition into euchromatin due to their deleterious effects for host cells that might be not only related to potential genetic alterations but also to epigenomic alteration via PEV. As shown in [131], the species-specific spreading ability of TEs is likely to be a driving force of their counter-selection. On the other hand, TEs are main contributors of genome evolution, by promoting genomic diversity and allowing regulatory innovations. Co-opted TEs play a crucial regulatory role in various nuclear processes, in particular in gene regulation during development [135, 136] as well as in 3D genome organisation [137]. TEs are constitutively silenced in most cell types and this silencing is required not only to limit their transposition but also to maintain proper cell-type 3D organisation and gene expression pattern. Hence, deregulation of TE has been associated to pathologies and is a clear hallmark of cancer [138, 139]. Overall this clearly indicates the need to develop a quantitative model based on the painter framework that would describe the epigenomic control of TEs and how they affect in turn the epigenome dynamics of flanking genomic regions. This will pave the way for a better understanding of the role of epigenome dynamics in TE-based genome plasticity during evolution as well as in pathologies.

## 5 ACKNOWLEDGEMENTS

We are grateful to Geneviève Fourel, Kapil Newar, Gaëlle Legube and Raphaël Mourad for fruitful discussions. We acknowledge Agence Nationale de la Recherche (DJ: ANR-18-CE12-0006-03; AZB, DJ: ANR-18-CE45-0022-01; DJ, CV: ANR-21-CE45-0011-01) for funding. We thank PSMN (Pôle Scientifique de Modélisation Numérique) and CBP (Centre Blaise Pascal) of the ENS de Lyon for computing resources.

