## Supplementary Information for "Painters in chromatin: a unified quantitative framework to systematically characterize epigenome regulation and memory"

### 1 Shapes of $P_c$

#### 1.1 Diffusion-limited case

To estimate $P_c(i,j)$ in the case where the effective spreading from position $i$ to a distal nucleosome $j$ occur by the unbinding and diffusing of the HME recruited in $i$, we follow a formalism consistent with Li et al. [1]. H. Berg [2] showed that the probability for a slowly-diffusing particle to hit an object of size $a$ initially at a distance $R$ from the particle is $P_d = a/R$. Within our context, $R$ is thus the typical distance between $i$ and $j$. In the main text, $R(i,j)$ is taken to be the root mean squared distance between two loci of a simple self-avoiding polymer $R(i,j) \sim s^{0.5}$ with $s = |j-i|$ the genomic distance between $i$ and $j$. This leads to $P_d(i,j) \propto 1/|j-i|^{0.5}$, that is why we take the corresponding spreading probability to be $P_c(i,j) = 1/|j-i|^{0.5}$.

#### 1.2 Loop extrusion case

To estimate $P_c$ in the case where spreading from a HME bound at $i$ to a distal nucleosome $j$ is possible only if they are placed in very close proximity by a loop extruding factor (LEF like cohesin or condensin) translocating along the chromatin, we develop a simple mathematical model (see Supplementary Fig. 2) [5–8]. We assume loop extruding factors are composed by two motor subunits [9] that bind to chromatin on adjacent sites $(i, i+1)$ with a fixed rate $k_{lb}$. Once bound, they translocate bidirectionally away from one another at constant speed $v$. LEFs may also unbind from the chromatin at a rate $k_{lu}$. For simplicity, we assume a low density of LEFs on chromatin and thus neglect collisions between them. We also neglect the presence of barriers (like CTCF) that may stop or slow down LEF subunit translocation [10]. From these hypotheses, the dynamics of the probability to find a LEF between $i$ and $j$ $(P_l(i,j))$ simply follows

$$\frac{dP_l(i,i+1)}{dt} = k_{lb} - (2v + k_{lu})P_l(i,i+1) \quad (1)$$

$$\frac{dP_l(i,j)}{dt} = v(P_l(i+1,j) + P_l(i,j-1)) - (2v + k_{lu})P_l(i,j) \quad (2)$$

At steady state, this leads to $P_l(i,i+1) = k_{lb}/(2v + k_{lu})$, and in general, for $j \neq i+1$, $P_l(i,j) = v(P_l(i+1,j) + P_l(i,j-1))/(2v + k_{lu})$. Assuming an homogeneous system, ie that $P_l(i,j)$ only depend on the genomic distance $s$ between $i$ and $j$ $(P_l(i,j) \equiv P_l(s)$ with $s = |j-i|)$, we obtain the recurrence relation

$$P_l(s) = \left(\frac{2v}{2v + k_{lu}}\right) P_l(s-1) \quad (3)$$

$$\text{with} \quad P_l(1) = \frac{k_{lb}}{2v + k_{lu}} \quad (4)$$

Therefore $P_l(s) = \beta^s \alpha$ with $\beta = 2v/(2v + k_{lu})$ and $\alpha = k_{lb}/(2v)$, leading to $P_l(s) = \alpha e^{-s/s_0}$ with $s_0 \equiv -1/\log\beta > 0$ a typical genomic distance that characterize the processivity of the LEF. For example, in the limit of fast extruding LEFs $(2v \ll k_{lu})$, $s_0 \approx (2v)/k_{lu}$. That is why we take the corresponding spreading probability to be $P_c(i,j) = e^{-s/s_0}$.

#### 2 Polymer Simulations

To estimate $P_c$ in the cases of a central, strongly-self-interacting domain of nine nucleosomes and of a long-range loop between the painter region and a distal region, we perform simulations of a simple, isolated self-avoiding polymer (composed by 201 beads of diameter 10 nm) using the lattice kinetic Monte-Carlo model developed in [4]. To simulate the single 3D compact domain (Fig. 5 lower part), we use a self-attraction of $-1kT$ between the nine nucleosomes surrounding the painter region. The same framework is used for the looping case (Fig. 5 upper part) with attractive interaction ($-1kT$) between monomers of the painter region (positions: $[-2 : 2]$) and monomers at positions $[48 : 52]$. To obtain the $P_c(i,j)$ matrix in each case, we simulate 128 independent trajectories by first letting the system to equilibrate during $9 * 10^7$ Monte-Carlo time Step (MCS) before taking measurements every $10^5$ MCS. From the ensemble of configurations, we thus estimate the contact probability $P_c(i,j)$ between any pairs of monomers as the probability that the relative distance between $i$ and $j$ is less than 40nm.

#### 3 Analytical solutions

In the case where state-specific recruitment is not present ($\Delta = 0$)

$$P(M_i) = \frac{(k/k_0)(\rho_s(i) + \varepsilon \sum_{j \neq i} \rho_s(j) * P_c(i,j)(1 + r\delta_{j,M}))}{(k/k_0)(\rho_s(i) + \varepsilon \sum_{j \neq i} \rho_s(j) * P_c(i,j)(1 + r\delta_{j,M})) + 1} \quad (5)$$

where $\delta_{j,M} = 1$ if nucleosome j is in M-state, $= 0$ otherwise. Inside the painter region, $\overline{P_p}$ (the average value of $P(M_i)$ inside the painter region) can be computed by assuming that $\delta_{j,M} = \overline{P_p}$. Under this mean)field approximation, $\overline{P_p}$ satisfies

$$\overline{P}_p = \frac{-(1 + c\varepsilon(1-r) + k_0/k)}{2\varepsilon rc} + \frac{\sqrt{(1 + c\varepsilon(1-r) + k_0/k)^2 + 4\varepsilon rc(1 + \varepsilon c)}}{2\varepsilon rc}. \quad (6)$$

where $c = \sum_{j \in painter} \rho_s(j) * P_c(i,j)$. For a painter region of size $N = 5$ nucleosomes, $c = 12.83$ if $P_c(i,j) = 1/|j - i|$. Outside the painter region, since $\rho_s(i) = 0$ (no sequence-specific recruitment), $P(M)$ follows Eq. 3 of the main text but by replacing $\varepsilon$ by $\varepsilon(1 + r\overline{P}_p)$ (triangles in Fig. 3). This solution can be improved in the range of high fluctuations (Fig. 3B main text), if we first compute $P(M_i)$ $\forall i \in painter$ from Eq. 5 by assuming that $\delta_{j,M} = P(M_i)$ and by solving the corresponding closed set of quadratic equations $\forall i \in painter$. Then, the average value of these solutions $\overline{P}_p^* = 1/L \sum_{i \in painter} P(M_i)$ is introduced in Eq. 5 (writing $\delta_{j,M} = \overline{P}_p^*$) to now derive the probability of M-state outside the painter region (circles in Fig. 3)

#### 4 Enzyme limitation

Here we consider explicitly that the M-state can recruit HMEs and thus may switch to an enzyme-bound state E, the transition rates being dependent on the binding-unbinding kinetics of the enzyme $k_b, k_u$ (Fig. 4A). Enzyme bound state has the ability to propagate modifications to neighbouring nucleosomes in 3D. Assuming fast binding-unbinding enzyme kinetics, we can simplify the formalism into a two - state system (consistent with the description in the article). Indeed, we can write $k_b N_e P(M) = k_u P(E)$ with $N_e = (N_e^{tot} - \sum_j \rho_s(j) - \sum_j \delta_{j,E})$ the current number of unbound enzymes where $N_e^{tot}$ is the total number of HMEs, $\sum_j \rho_s(j)$ is the number of sequence-specifically bound enzymes and $\sum_j \delta_{j,E}$ the number of HMEs bound to M-states. By approximating $\sum_j \delta_{j,E} \approx \left[\sum_j \delta_{j,M}\right] P(E)$ the following expression (Eq. 7) is obtained for $P(E)$:

$$P(E) = \left[2\sum_j \delta_{j,M}/(N_e^{tot} - \sum_j \rho_s(j))\right]^{-1} \left[1 + \frac{\sum_j \delta_{j,M}}{N_e^{tot} - \sum_j \rho_s(j)} + \frac{k_u}{k_b(N_e^{tot} - \sum_j \rho_s(j))} - ((1 + \frac{\sum_j \delta_{j,M}}{N_e^{tot} - \sum_j \rho_s(j)} + \frac{k_u}{k_b(N_e^{tot} - \sum_j \rho_s(j))})^2 - \frac{4N_M}{N_e^{tot} - \sum_j \rho_s(j)})^{1/2}\right] \tag{7}$$

$P(E)$ is then used to re-normalize the state-dependent term in Eq. 1 (main text) to have accounted for enzyme limitation in the model. In this case, the modified state represents a lump state consisting of the M-state and the enzyme bound state $E$ and the spreading process depends on $N^{tot}$ and $k_b/k_u$. For a constant $k_b/k_u = 1$, $P(M)$ in a region is dependent on the total number of HMEs. In the presence of sequence-dependent recruitment, enzyme limitation can confine the domains around the painter region (Fig. 4B) while in the absence of sequence dependent recruitment, we have a uniform steady state profile of M-state whose absolute value is dependent on $N_e^{tot}$ (Fig. 4C,D).

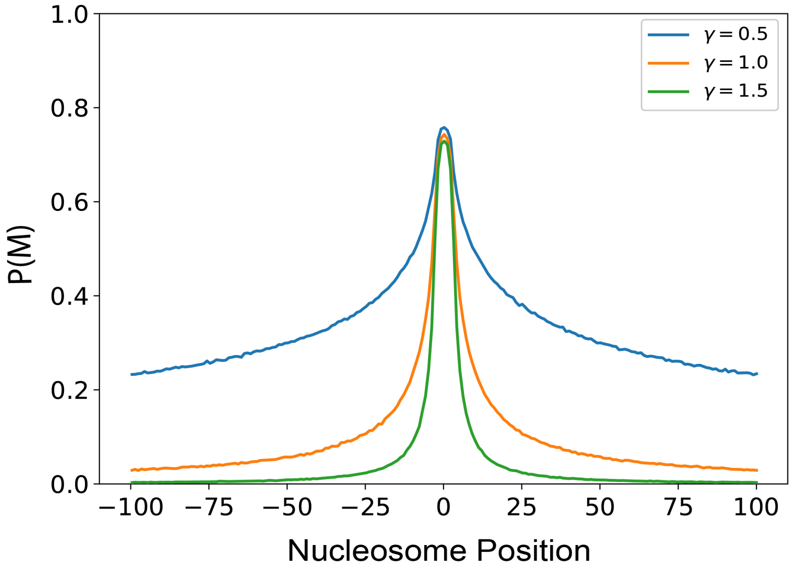


**Supplementary Figure 1.** Virtual Chip-seq profiles of the M-state as a function of the exponent $\gamma$. We show how chromatin compaction affects the process of spreading of M-state. A compact chromatin region means low $\gamma$-values.

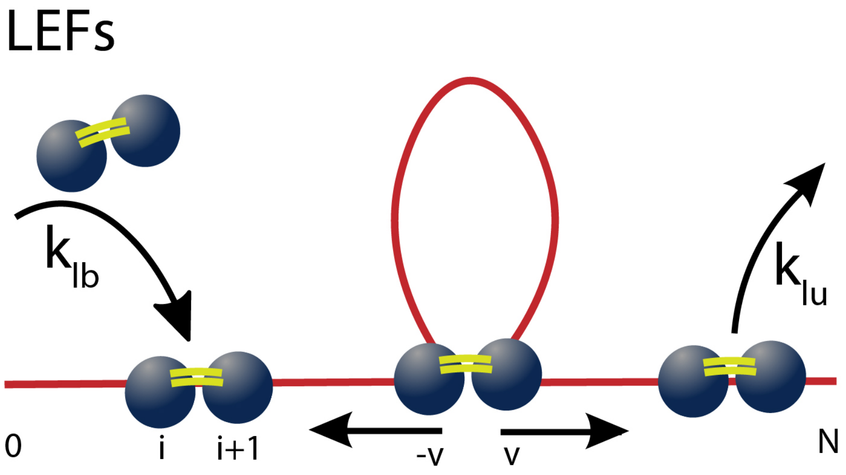


**Supplementary Figure 2.** Illustration of the loop extrusion model. Loop extruding factors (LEFs) can bind to the chromatin (in red) at position $i, i+1$ with a fixed rate $k_{lb}$ and unbind at a rate $k_{lu}$. The two subunits of the LEFs move bidirectionally in opposite directions at velocity $v$ leading effective extrusion of chromatin loops.

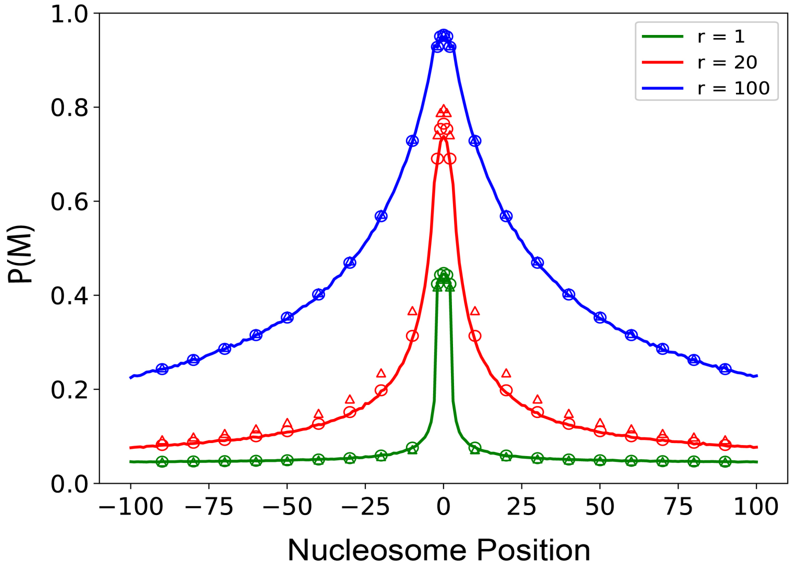


**Supplementary Figure 3.** Comparison between the analytical solutions and simulations in the boosted-painter mode. The different $r$ values correspond to stable ($r = 100$) and highly fluctuating ($r = 20$) situations in Fig. 3B (main text). The triangles are analytical solutions obtained by using $\overline{P}_p$ (from Eq. 6). The circles are obtained by using $\overline{P}_p^*$ to compute $P(M_i)$ outside the painter region which works better compared to the previous method.

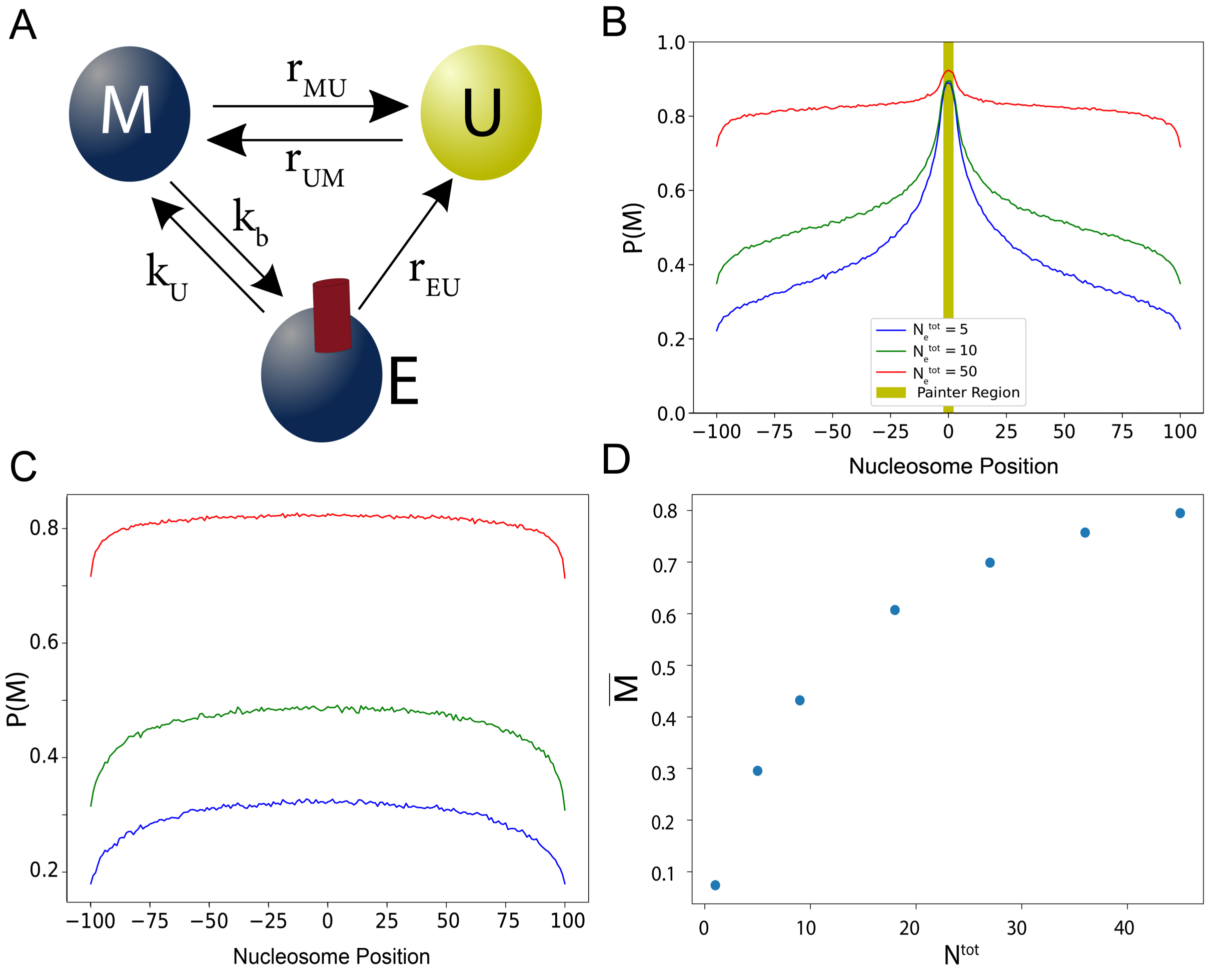


**Supplementary Figure 4.** Enzyme limitation: (A) Scheme of the enzyme limitation model. A modified nucleosome (M) can be in one of the two states: HME-free and HME-bound (E). The switching rate between these states is driven by the HME binding rate $k_b$ and the corresponding unbinding rate $k_u$. We assume that the propensity to switch to the unmodified state $U$ is similar in $M$ or $E$ states ($r_{EU} = r_{MU}$). A U-state is assumed to be HME-free. (B) Virtual Chip-seq profiles of the M-state (HME bound+HME-free modified state) with sequence-dependent recruitment at the painter region for different total number of HMEs $N_e^{tot}$ with $k/k_0 = 2$, $\varepsilon = 0.9$ and $k_u/k_b = 1$. $P_c(s) = 1/s$. (C) The same profiles as in (B) but in absence of sequence-dependent recruitment, showing the unconfined uniform spreading of the M-state for all $N^{tot}$ values. $P_c(s) = 1/s$. (D) Variation of $\bar{M}$ as a function of $N_e^{tot}$ in absence of sequence-dependent recruitment.

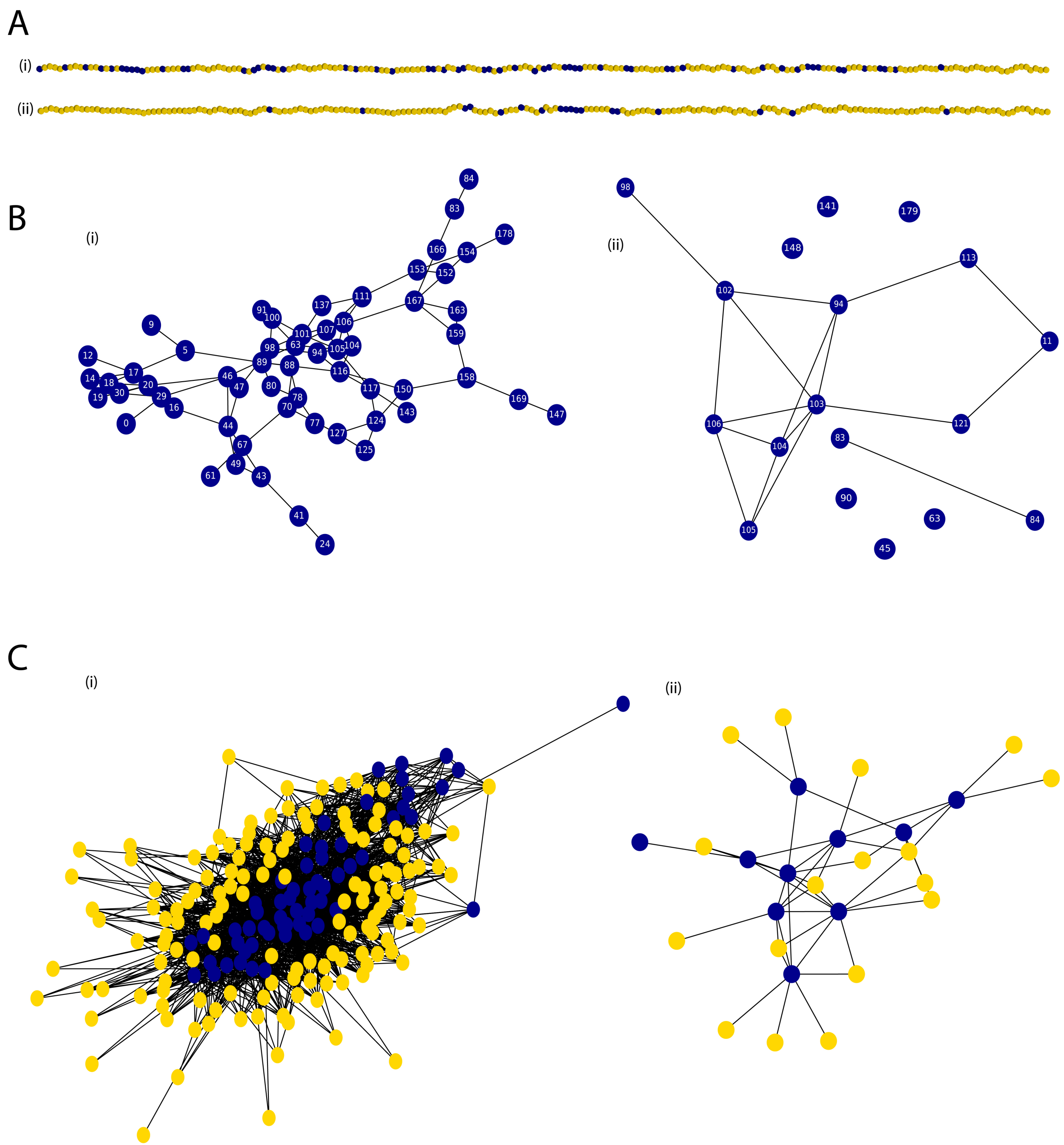


**Supplementary Figure 5.** (A) Two examples of the epigenomic state inside the region of interest (blue: M-state; yellow: U-state). Simulations were performed with $k/k_0 = 1$, $\varepsilon = 0.6$, $P_c(i,j) = 1/|i-j|$, and (i) $\Delta = 0.2$ and (ii) $\Delta = 0.0$. (B) For each example in (A), we plot one possible realization of the contact graph ($M_{net}$) (see main text). In (i), all the M-state nodes are connected inside one subgraph of size 59. In (ii), there are multiple disconnected M-state subgraphs, the largest one being of size 10. (C) For the largest subgraph show in (B), we plot one possible realization of an extended domain $M_{net}*$(see main text) including U-state nucleosomes. The blue nodes form the (modified) contact domain and the yellow nodes are unmodified nucleosomes in close proximity in 3D to the nucleosomes in largest M-state network $M_{net}(*)$. The extended domain size (total number of nodes) is (i) 198 and (ii) 27 in the two different cases.

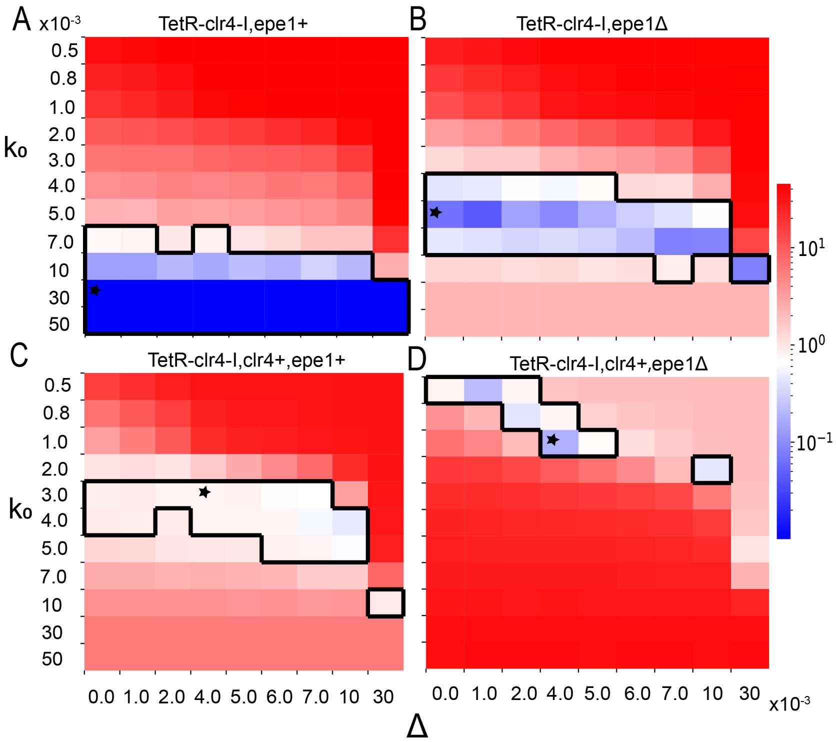


**Supplementary Figure 6.** Phase diagram showing the $\chi^2$ value that quantifies the predictive power of the model predictions relatively to the experimental data for each of the four mutants (see main text). The region within the highlighted contour represents the region of the parameter space with an acceptable fit ($\chi^2 < 1$). Star marks the best fitting parameters with respect to the experimental constraints. These parameters are used in Fig. 8C,D (main text).

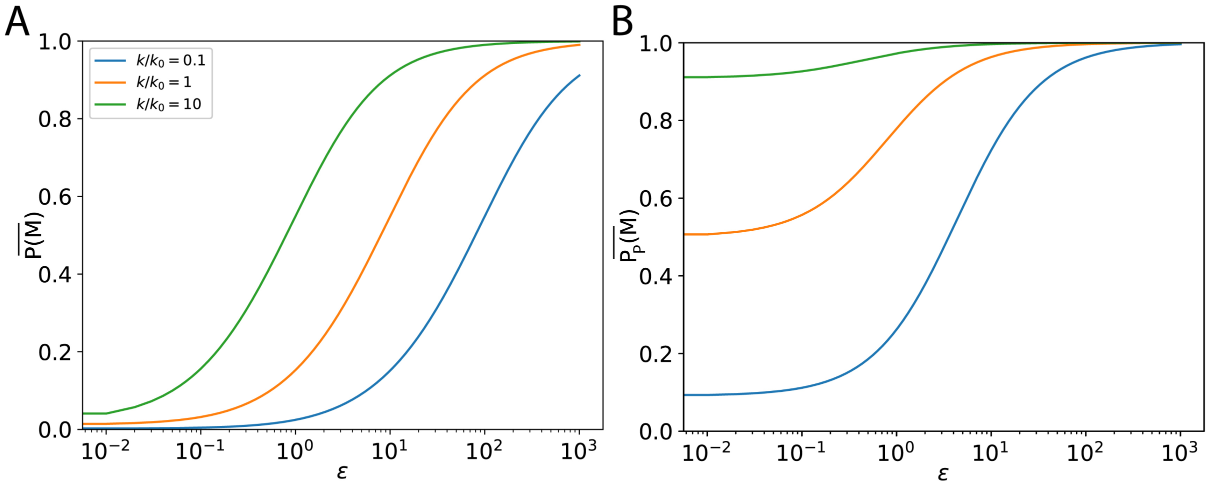


**Supplementary Figure 7.** Painter mode: the transition from low to high M-state as a function of the spreading efficiency ($\epsilon$) for different $k/k_0$ and $P_c(s) = 1/s$. (A) Global average of $P(M)$. (B) Corresponding average value of $P(M)$ within the painter region.

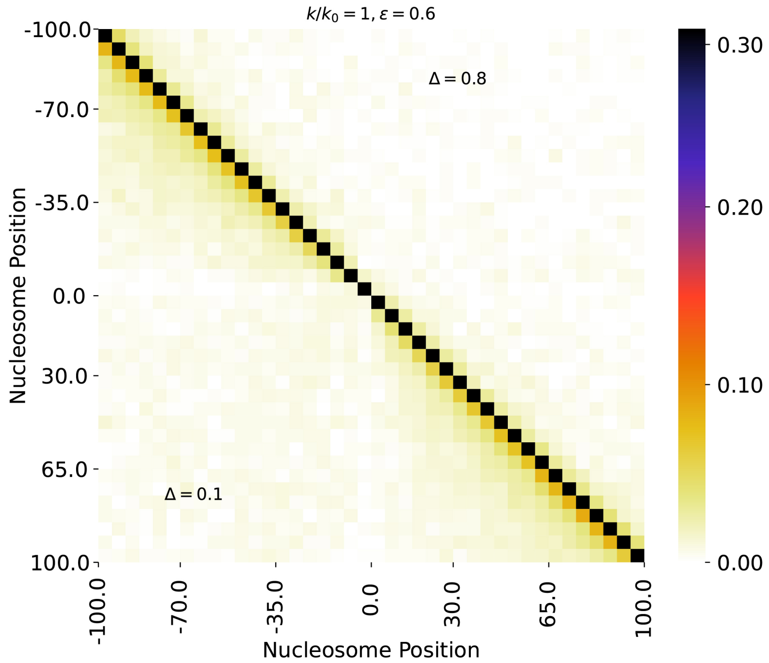


**Supplementary Figure 8.** Correlation matrices between the nucleosome state at two positions along the genomic region in the reader-writer mode, for $\varepsilon = 0.6$, $k/k_0 = 1$ and $\Delta = 0.1$ (lower part) and $\Delta = 0.8$ (upper part), $P_c = 1/s$. Reduced correlations are observed when $\Delta$ is far from the critical value ($\Delta_c \approx 0.2$ for such $\varepsilon$, $k/k_0$ values) (see also Fig. 4C main text).

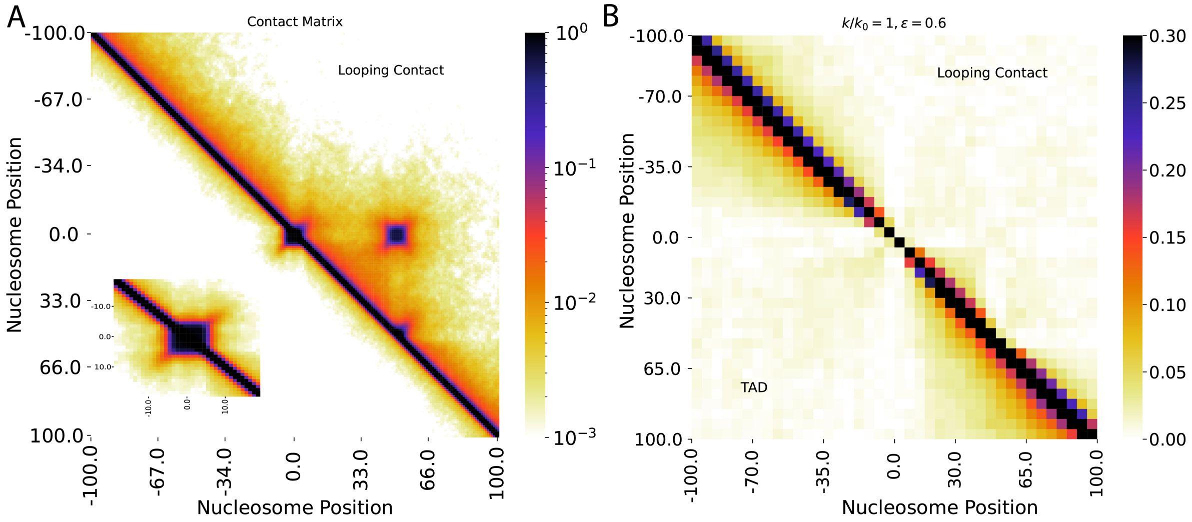


**Supplementary Figure 9.** (A) Contact probability matrix for a compact domain (lower part) localized at positions [-4:4] (inset shows a zoom around the compact domain) and looping contact (upper part) between the painter region [-2:2] and nucleosomes at position [48:52]. (B) Nucleosome state correlation in the compact domain (lower part) and looping (upper part) cases in the reader-writer mode (parameters:$k/k_0 = 1$, $\epsilon = 0.6$) at the critical value $\Delta_c = 1.2$ for contact domain and $\Delta = 0.3$ for the loop case.

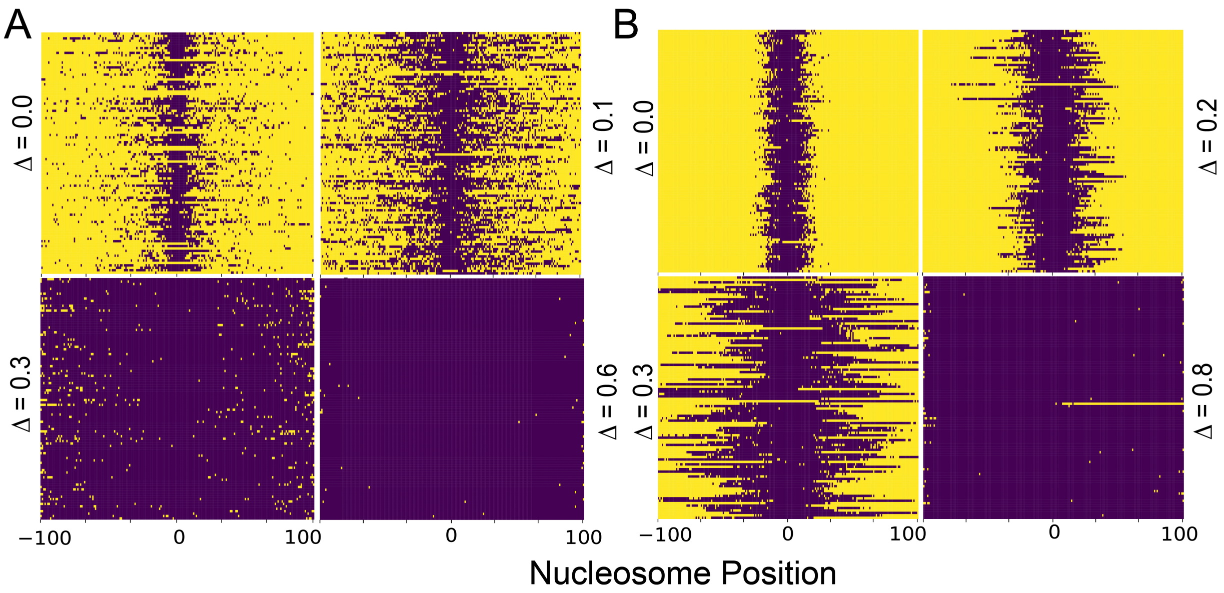


**Supplementary Figure 10.** (A) Illustration of the percolation of the system. For four different $\Delta$ values in 3D contact spreading probability case ($P_c(s) = 1/s$, orange curves in Fig.5B,C), the most extended generalized domains observed in different configurations (100) are stacked together to show the system stochasticity (all nucleosomes belonging to an extended domain are colored in blue). At high enough $\Delta$ ($= 0.6$), the system is fully percolated. (B) As in (A) but for the loop extrusion spreading mechanism.

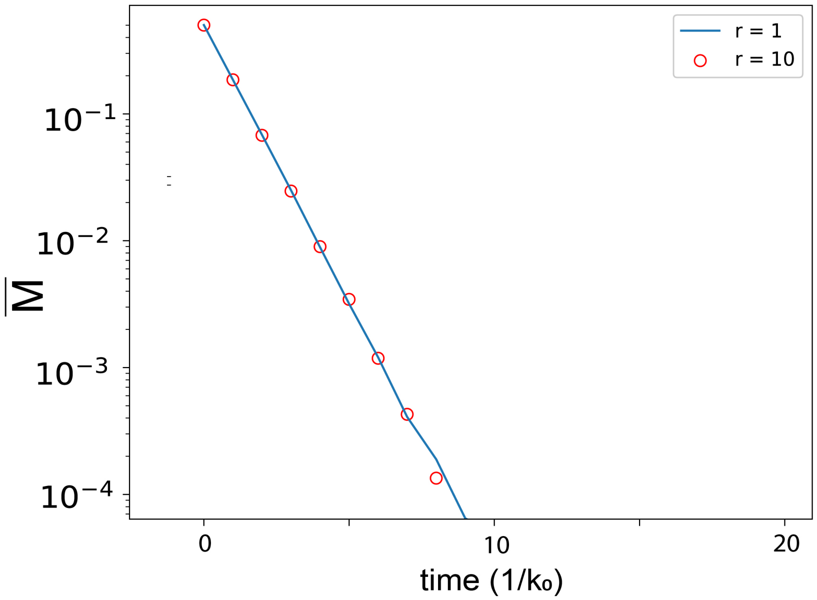


**Supplementary Figure 11.** Memory in the booster-painter mode. To investigate epigenetic memory, we study the evolution of $\overline{M}$ after the sequence dependent HMEs unbind at $t = 0$. Even though boosted-painter mode has state-dependence within the painter region, once the sequence-dependent HMEs unbind there is no more cooperativity hence, the decay of $\overline{M}$ is the same for different boost factor $r$ values unlike in the reader-writer mode where there is a strong dependency in the parameter $\Delta$ (see Fig. 5 main text).

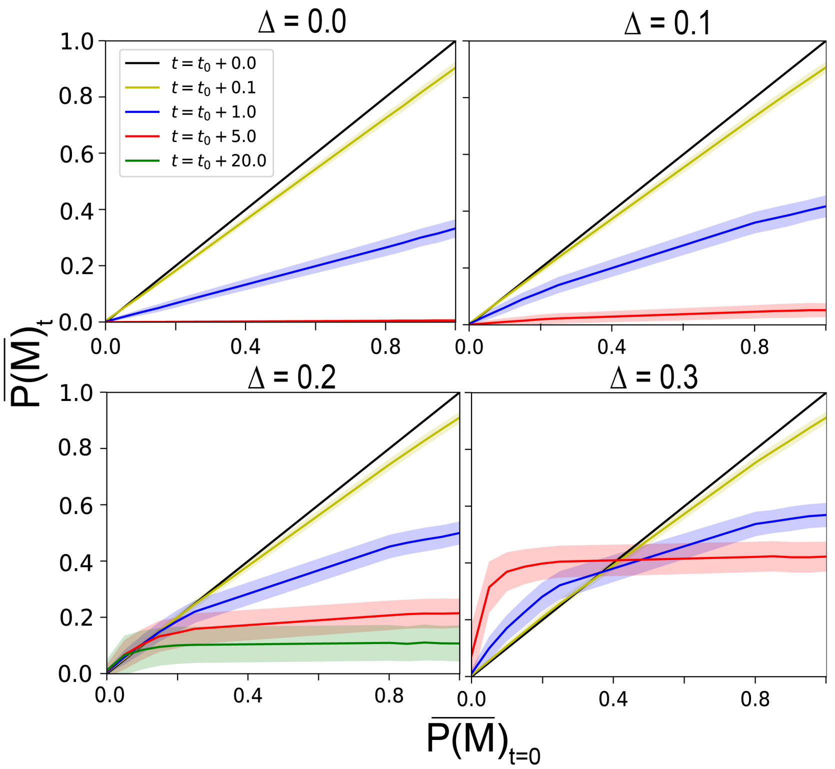


**Supplementary Figure 12.** To investigate the correlation between the M-state maintained by reader-writer mechanism at time $t$ and the initial state at time $t = 0$ just before the sequence-dependent painters unbind, we estimate $\overline{P(M)}_t$, the probability to be modified at a time $t$ after the unbinding of painters, as a function of an imposed initial state with fixed $\overline{P(M)}_{t=0}$. An $x = y$ scenario means perfect correlation (when the exact state is maintained). For $\Delta = 0.3$, even though the M-state is maintained it is not correlated with the state at $t = 0$. In general, time correlation of M-state is quickly lost which means the reader-writer mechanism would be able to maintain the level of repression but not exactly the shape of the profile.

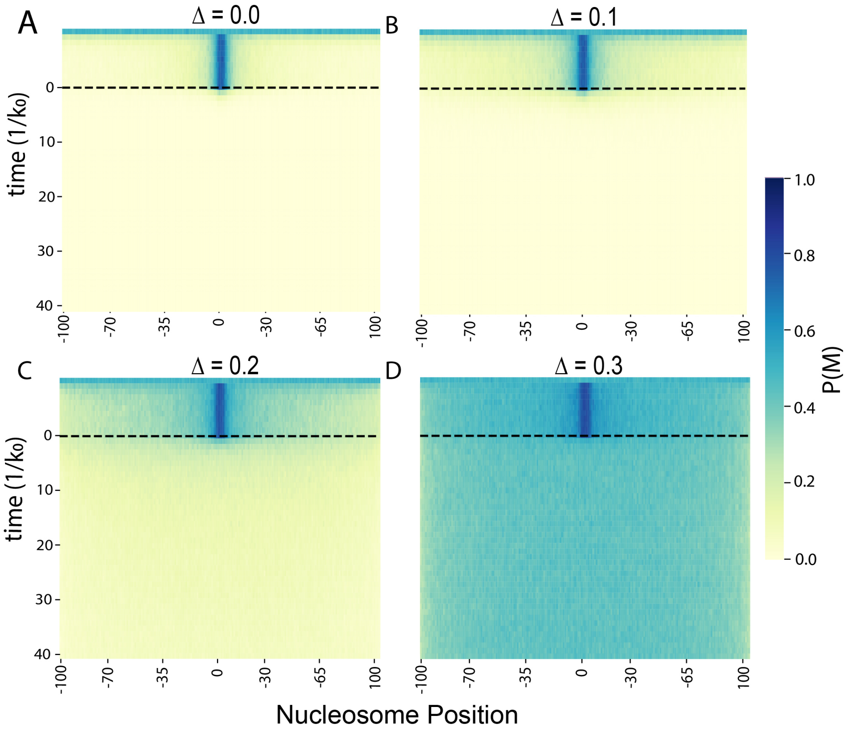


**Supplementary Figure 13.** Time evolution of $P(M)$ after the unbinding of the sequence-dependent painters at $t = 0$ (dotted black lines) for different values of $\Delta$ with $k/k_0 = 1$ and $\varepsilon = 0.6$ as shown in Fig. 5A inset of main text. Here $\forall\Delta$, the final state is a uniform state with no sequence-dependency.

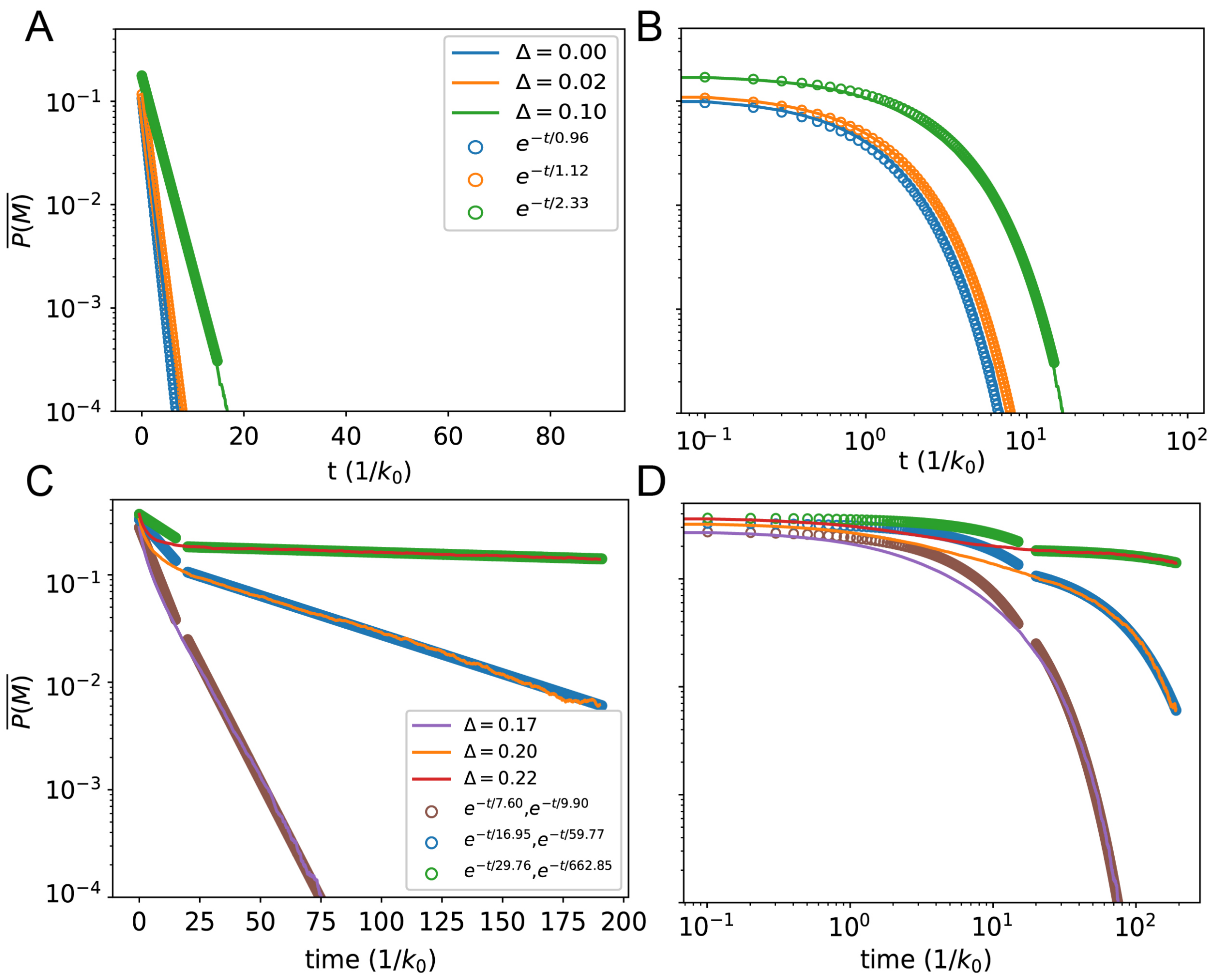


**Supplementary Figure 14.** Decay of the average M-state value in the region shown for different $\Delta$ values in log-lin scale (A,C) or log-log scale (B,D). Lines represent simulation results, dots correspond to the fit. (A,B) For $\Delta << \Delta_c = 0.2$, the decay is purely exponential. (B) (A) in log-log scale. (C,D) For $\Delta \leqslant \Delta_c$ the decay can be fitted by two time scale exponentials: a fast exponential decay at small time for $t \ll t*$ ( $\overline{P(M)} = D_0 e^{-t/\tau_0}$) and a much slower exponential at large time for $t \gg t*$ ($\overline{P(M)} = D_1 e^{-t/\tau_1}$ with $\tau_1 > \tau_0$). Exact fitted values are shown in the plot.

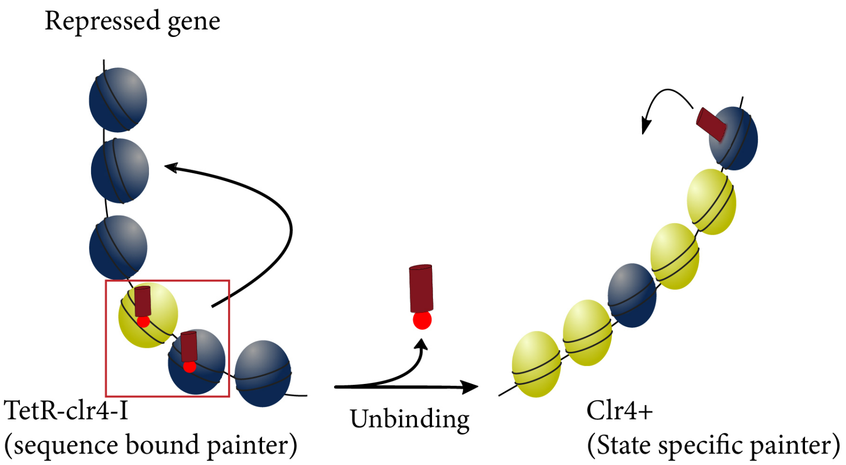


**Supplementary Figure 15.** Scheme describing the experiment on engineered yeast to study heterochromatin memory (Fig. 9C,D). Initially, the HMEs recruited at the target region establish repression (repressed gene state; (blue) modified nucleosomes) and then subsequently, once the HME unbind, the gene expression is monitored over a period of 100 hours in the four different mutants (see main text, Fig. 9C,D). After the sequence-dependent painter unbinds, all mutants loose gradually repression but at different rate (Fig. 9D), as the level of methylated nucleosomes decreases. In some mutants where *Clr4+* is present (red and magenta, Fig. 9D), after removing the sequence-specific recruitment, the reader-writer mechanism fights against this loss of repression by maintaining for some time a methylated state modifications.
